# Centromeres are Hotspots for Chromosomal Inversions and Breeding Traits in Mango

**DOI:** 10.1101/2024.05.09.593432

**Authors:** Melanie J. Wilkinson, Kathleen McLay, David Kainer, Cassandra Elphinstone, Natalie L. Dillon, Matthew Webb, Upendra K. Wijesundara, Asjad Ali, Ian S.E. Bally, Norman Munyengwa, Agnelo Furtado, Robert J. Henry, Craig M. Hardner, Daniel Ortiz-Barrientos

**Author notes:** Melanie J. Wilkinson, **Email:**. **Author Contributions:** M.J.W. and D.O. designed the study. N.D., A.A. and I.B. collected the data. N.D., A.A., I.B., M.J.W., N.M., U.K.W., A.F., R.J.H. and C.M.H. curated the data. M.W. generated alignments and variant calling. M.J.W. performed quality filtering on genotype data, LD, population genomic, GWAS and deleterious score analyses. C.E. predicted centromere locations. K.M. performed local PCA analysis. U.K.W., A.F. and R.J.H. completed HiFi genome comparisons. D.K. performed the joint contribution of inversions on trait value and enrichment analyses. D.O. and C.H. secured funding and were mentors. M.J.W. and D.O. wrote the manuscript with contributions from all authors.

## Abstract

Chromosomal inversions can preserve combinations of favorable alleles by suppressing recombination. Simultaneously, they reduce the effectiveness of purifying selection enabling deleterious alleles to accumulate. This study explores how areas of low recombination, including centromeric regions and chromosomal inversions, contribute to the accumulation of deleterious and favorable loci in 225 *Mangifera indica* genomes from the Australian Mango Breeding Program. Here, we identify 17 chromosomal inversions that cover 7.7% (29.7Mb) of the *M. indica* genome: eight pericentric (inversion includes the centromere) and nine paracentric (inversion is on one arm of the chromosome). Our results show that these large pericentric inversions are accumulating deleterious loci, while the paracentric inversions show deleterious levels above and below the genome wide average. We find that despite their deleterious load, chromosomal inversions contain small effect loci linked to variation in crucial breeding traits, indicating that chromosomal inversions have likely facilitated their selection. The results from this study have important implications for selective breeding of favorable combinations of alleles in regions of low recombination.

**Significance Statement:** Chromosomal inversions and other low recombination regions of the genome can drive trait evolution. Fewer recombination events can assist in maintaining favorable combinations of alleles, but it can also make disentangling favorable and deleterious alleles difficult. Understanding whether these low recombination regions contain favorable or deleterious loci could drive our decision to increase or decrease the frequency of these regions in target breeding populations. Breeding for large segments of the genome based on presence or absence of an inversion can rapidly drive large trait differences within few generations. Harnessing the impact of large low recombination regions of the genome could have major implications for future genetic improvement in breeding.

## Introduction

Low recombination regions of the genome, such as those harboring chromosomal inversions, play a major role in trait evolution. An inversion occurs when a segment of chromosome breaks off and re-attaches in the reverse direction. Chromosomal inversions are prevalent in natural (1–5) and domesticated plant populations (e.g. rice (6, 7), wheat (8), barley (9), cucumber (10), melon (11), soybeans (12), cotton (13) and tomatoes (14)). The mismatch of loci between the inverted and non-inverted segments limits recombination between them, enabling inversions to maintain combinations of alleles (15). The accumulation of selectively advantageous alleles in inverted segments highlights their evolutionary role in trait evolution. For example, inversions have been implicated in adaptive evolution across a wide-range of species, including sunflower (3), monkeyflowers (5), threespine sticklebacks (2), cod (4), and deer mice (1). Despite the known role of inversions in driving the evolution of favorable traits in nature, selecting for desirable inversions rarely occurs in plant breeding populations (16).

Selective breeding for inversion variation has often occurred incidentally during domestication. When the genetic architecture of these traits is largely captured by inversions, selection for these traits drives the frequency of the inversion to increase in the population. For example, selection for domestication genes in soybean likely drove a sweep for a large inversion (43.30-43.66Mb) on chromosome seven (12). In rice, inversions on chromosome six contain genes for rice blast disease resistance (7) and in *Brassica napus* researchers found inversions persistently contained disease resistance genes (17). It is often difficult to assess whether the inversion or the desirable alleles arose first, but evidence of inversions harboring desirable breeding traits continues to emerge as researchers identify chromosomal inversions in their breeding populations (16). Harnessing these low recombination regions, containing linked desirable combinations of alleles, for selective breeding could open the door for simple genotyping of inversion presence/absence and enable substantial changes in multiple traits in few generations.

Methods for identifying chromosomal inversions often require long read sequencing on few individuals. However, this is an expensive approach, so methods that can quickly and cheaply scan the genome for candidate inversions are a great complement to study regions of low recombination. Local principal component analysis (PCA), which uses inexpensive short read sequencing, can reveal candidate chromosomal inversions that are driving genetic differentiation (3, 18). The local PCA approach identifies outlier regions that are more genetically structured than the genome average. A typical signature of an inversion segregates these outlier regions into three clusters, predicted to reflect 0, 1 or 2 copies of the inversion for diploid species. If heterozygosity is high in the group of individuals predicted to carry one copy of the inverted chromosomal segment, we could more confidently infer the presence of an inversion.

Furthermore, genetic differentiation should be maximum between the two homozygous clusters, which reflect groups of individuals with collinear chromosomes (i.e., 0 or 2 copies of the inferred inversion). Because recombination is prevented between the homozygous clusters, we also expect to see a drop in nucleotide diversity and an increase in linkage disequilibrium in these potential inversion regions. This chromosomal inversion identification approach will likely identify large inversions driving population differentiation, which could uncover the variation underlying key traits for selective breeding.

Low recombination regions of the genome accumulate deleterious alleles in natural and domesticated populations (19–23). Fewer recombination events during meiosis in regions containing chromosomal inversions and centromeres, prevent the effective separation and removal of mutations that reduce fitness from those that increase fitness (24). New mutations arise across the genome every generation, and deleterious mutations arise more frequently than beneficial mutations, as there are more ways to disrupt function than improve function (25). As such, regions of low recombination can become sinkholes for mutations with slight to moderate deleterious effects on populations. Gossmann*, et al.* (26) found that on average in plants, 25% of new nonsynonymous mutations (those that change the amino acid) behave as effectively neutral, the remaining being mostly deleterious with few being advantageous. Factors, such as selection (natural or artificial) and lack of genetic diversity, can amplify the accumulation of deleterious mutations in regions of low recombination (22, 27). One example includes selection for favorable alleles that leads to an increase in frequency of harmful alleles that are in genetic linkage (28).

Another factor that is prevalent in breeding populations is low genetic diversity from recent bottlenecks, which can drive slightly deleterious alleles to become fixed in the population (29). If there are no other haplotypes without the deleterious alleles, then there is no opportunity for selection to purge the harmful alleles until new variation is introduced into the population. These deleterious alleles are thought to cause inbreeding depression (30), which can drive reductions in yield. Thus, crop breeding could be improved if deleterious mutations are removed from the genome (22).

Here we explore how regions of low recombination in the genome - centromeres and chromosomal inversions - contribute to the accumulation of desirable and deleterious loci in 225 *Mangifera indica* genomes from the Australian Mango Breeding Program. We identified candidate centromere locations, using tandem repeat signatures in the *M. indica* cv. ‘Alphonso’ genome.

The local PCA approach was implemented to identify candidate chromosomal inversions across the population, and these results were corroborated using long read sequencing on few individuals. To identify desirable loci, we found sites underlying five key mango breeding traits – fruit blush color, fruit firmness, fruit weight, total soluble solids (proxy for sweetness) and trunk circumference (proxy for tree size). To identify deleterious loci we implemented a sequence homology approach (Sorting Intolerant From Tolerant For Genomes (31, 32)), where non-synonymous mutations in regions conserved across many species will produce higher deleterious scores. Our study seeks to understand the impact of low recombination regions of the genome on the balance between the accumulation of desirable and deleterious alleles in breeding populations, which has important implications for simplifying breeding practices.

## Results

### Centromeres accumulate genetic differentiation and chromosomal inversions in *M. indica*

To identify areas of genetic differentiation in the mango gene pool, we performed a local PCA, as implemented in Huang et al. (3), on SNPs derived from whole genome sequences of 225 *Mangifera indica*. We found 28 genomic regions (0.16 - 4.43Mb in size) across 17 chromosomes (total of 20; SI Appendix, Table S1) displaying extreme genetic differentiation (4 SD from the genome-wide average; SI Appendix, Fig. S1). We estimated the centromere location in the *M. indica* cv. ‘Alphonso’ genome using tandem repeats (±1Mb; SI Appendix, Fig. S2 and Table S1) (33). We discovered half of the regions of major differentiation overlap the predicted centromere (SI Appendix, Table S1), indicating that low recombination regions of the genome might play a role in allowing genetic differentiation in *M. indica* to persist.

We tested for differences in homozygosity between local PCA clusters to identify potential chromosomal inversions. For each region of major differentiation identified by local PCA, individuals were separated into three genotype groups using kmeans clustering. The central cluster is expected to be heterozygous for the inversion and the other two are predicted to contain alternate homozygote individuals for the inversion. Thus, if measured homozygosity significantly differs between the predicted heterozygous cluster and the two predicted homozygous clusters, we will consider these regions as potential chromosomal inversions. Homozygosity was differentiated in the expected direction in 17/28 genomic regions displaying extreme genetic differentiation (SI Appendix, Fig. S3). Thus, we further investigated these 17 potential chromosomal inversions across the *M. indica* genome (Table 1), where eight were pericentric (inversion contains the centromere) and nine paracentric (inversion is on one arm of the chromosome). These 17 potential chromosomal inversions, encompassing 7.7% of the genome, were abundant around centromeres, which further decreases the recombination events in this area of the genome.

**Table 1.** The 17 putative chromosomal inversions (inv) and their locations in the *M. indica* cv. ‘Alphonso’ (CATAS_Mindica_2.1) genome. Inversions are named according to the order to which they appear on a chromosome e.g. the first *M. indica* inversion on chromosome 1 (miinv1.0). Inversions are pericentric (located across the centromere) or paracentric (located on one arm of the chromosome). Sixteen of the inversions identified using a local PCA were corroborated with comparisons between the ‘Alphonso’ genome and haplotypes of three high fidelity (HiFi) genomes. The number of HiFi inversions identified within each of the local PCA inversions is shown. These inversions have a deleterious score lower (-) or higher (+) than the genome average (quantiles 2.5% = 0.483 and 97.5% = 0.487 from 1,000 bootstraps). Deleterious scores are represented as + >0.487, ++ >0.524, +++ >0.560, - <0.483, - - <0.443 and, - - - <0.402. Trait associations with each inversion were tested using two methods, an iterative random forest (ranking 1-3 shown) and a linear model (*P ≤ 0.05, **P ≤ 0.01, ***P ≤ 0.001). The five mango traits: fruit blush color (BC), fruit firmness (FF), fruit weight (FW), total soluble solids (TSS) and trunk circumference (TC).

| Inv | Inv type | Inv start (Mb) | Inv end (Mb) | Region size (Mb) | N HiFi invs within region | Deleterious score | Trait associations |  |  |  |  |
| --- | --- | --- | --- | --- | --- | --- | --- | --- | --- | --- | --- |
|  |  |  |  |  |  |  | BC | FF | FW | TSS | TC |
| miinv1.0 | Pericentric | 12.3 | 15.8 | 3.5 | 4 | ++ | ** |  |  |  |  |
| miinv3.0 | Pericentric | 14.6 | 17.9 | 3.3 | 6 | + | * |  |  | 1 |  |
| miinv4.1 | Pericentric | 8.3 | 9.3 | 1.0 | 0 | - |  | 1** |  | 3** |  |
| miinv5.0 | Pericentric | 3.9 | 7.4 | 3.6 | 1 | ++ |  | 2 |  |  | 2 |
| miinv8.0 | Pericentric | 14.3 | 18.2 | 4.0 | 5 | ++ |  |  |  |  |  |
| miinv9.0 | Pericentric | 5.1 | 7.0 | 1.9 | 2 | - | 2* |  |  |  |  |
| miinv14.0 | Pericentric | 8.7 | 11.5 | 2.8 | 4 | - |  |  |  |  |  |
| miinv20.0 | Pericentric | 4.1 | 5.3 | 1.2 | 3 | --- |  | * | *** |  |  |
| miinv1.1 | Paracentric | 16.3 | 16.5 | 0.2 | 1 | --- | 3 | 3 | 2 | * |  |
| miinv1.2 | Paracentric | 16.9 | 17.3 | 0.4 | 1 | --- |  |  |  |  |  |
| miinv4.0 | Paracentric | 5.4 | 7.0 | 1.6 | 5 | +++ |  |  |  |  |  |
| miinv6.0 | Paracentric | 9.6 | 10.1 | 0.5 | 1 | + | 1*** |  | 1*** | 2 |  |
| miinv7.0 | Paracentric | 15.4 | 17.9 | 2.5 | 1 | - |  |  | 3*** |  |  |
| miinv11.0 | Paracentric | 15.5 | 16.5 | 1.0 | 1 | + |  |  |  |  | 3 |
| miinv13.0 | Paracentric | 6.3 | 7.5 | 1.2 | 1 | ++ |  |  |  |  |  |
| miinv17.0 | Paracentric | 0.8 | 2.1 | 1.3 | 1 | - |  | * |  |  | 1*** |
| miinv17.1 | Paracentric | 1.3 | 2.0 | 0.6 | 1 | - |  |  |  |  |  |

If these 17 potential inversions identified by the local PCA are true inversions, we expect to see specific signatures in population genomic analyses (Fig. 1). All 17 potential inversion regions often cluster into three distinct groups in multi-variate space on PC1 (SI Appendix, Fig. S4), with the discreteness of the clusters ranging between 0.85-0.98 (maximum discreteness is 1; SI Appendix, Table S3). Additionally, genetic differentiation (FST) between the homozygous clusters (clusters 0 and 2) was 4.9-fold higher within the 17 potential inversions compared to the rest of the genome (FST Mean ± SD: inv = 0.49 ± 0.22 and non-inv = 0.10 ± 0.12), consistent with these clusters being fixed for opposing haplotypes. The smaller *M. indica* inversion on chromosome 17, miinv17.0, was the only candidate not showing signs of elevated FST, which might have been driven by miinv17.1, which overlaps this region. Pericentric inversion regions exhibited higher genetic differentiation compared to paracentric (FST: pericentric = 0.56 ± 0.20 and paracentric = 0.34 ± 0.17), with pericentric regions containing the highest FST peaks. There was a reduction in nucleotide diversity in at least one of the homozygous clusters in most (7/8) pericentric inversion regions (SI Appendix, Fig. S3), as expected for areas of reduced recombination. On the other hand, only 3/9 paracentric inversions showed a dip in nucleotide diversity within the inversion region (SI Appendix, Fig. S3), suggesting greater gene exchange (known as gene flux (34)) between clusters in paracentric inversions. Overall, our findings highlight that pericentric inversions in *M. indica* have high genetic differentiation and low nucleotide diversity, consistent with chromosomal inversions that reduce the exchange of alleles between chromosomes that contain the inversion and those that do not. On the other hand, paracentric inversions have less distinct inversion signatures, as they seem to have experienced some gene flux between clusters.

**Fig. 1.**
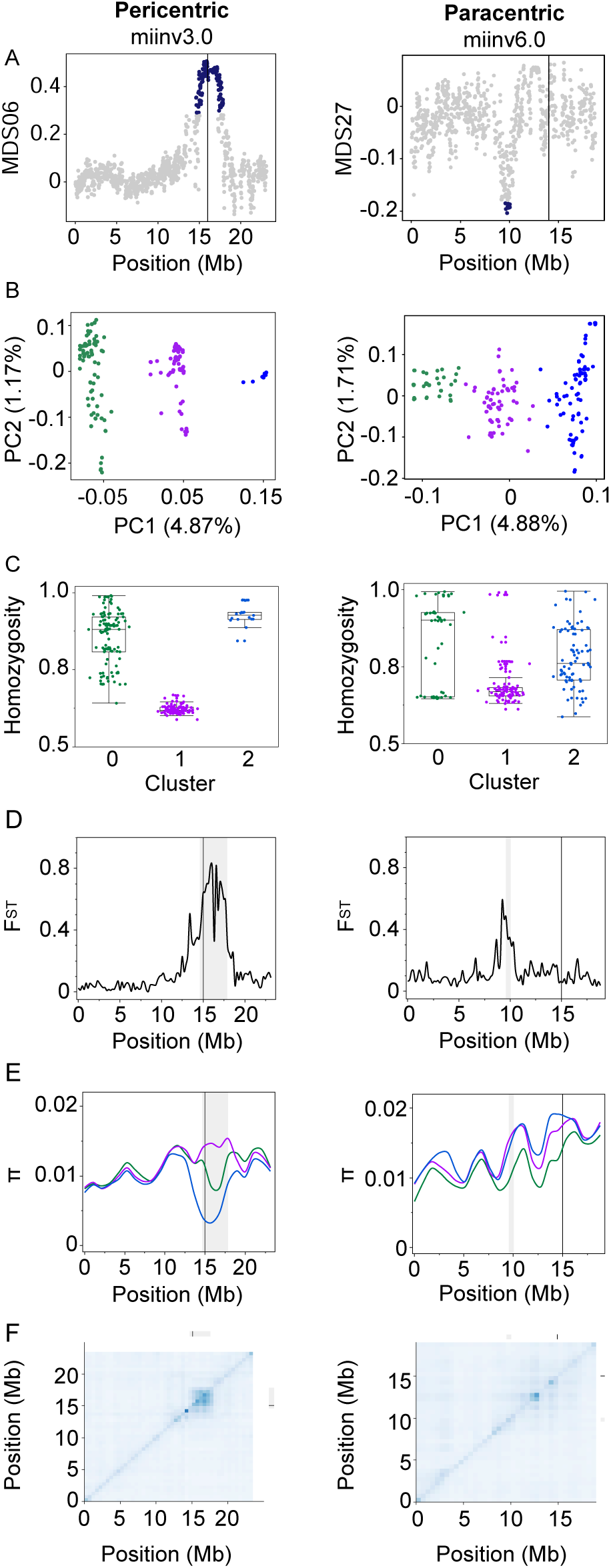
Identifying putative chromosomal inversions using a local principal component analysis (PCA). A) Local PCA multidimensional scaling plots, where the dark dots show the outlier region. B) PCA of the outlier region. C) Homozygosity of the outlier region for the three clusters. D) Genetic differentiation (F_ST_) between homozygous clusters, 0 and 2 across the chromosome. E) Nucleotide diversity (π) for the three clusters. F) Linkage disequilibrium heatmap of the most common homozygous group on the top triangle and all individuals on the bottom triangle (including the middle line). The black line is the predicted centromere, and the grey bar is the predicted inversion region. The colors represent the k-mean clustering (cluster 0 = green, cluster 1 = purple, and cluster 2 = blue).

To assess whether the predicted centromere and inversion regions have signatures of reduced recombination, we evaluated linkage disequilibrium (LD) patterns. Consistent with few recombination events, there was high LD around the centromere in all chromosomes that could be assessed (n=19), revealing the role the centromere plays in reducing recombination in this gene pool. As expected, most potential inversions show an increase in LD across all genotypes in the inversion region compared to the surrounding genomic regions (SI Appendix, Fig. S3). For paracentric inversions, genotypes with the same chromosomal arrangement (same cluster) should recombine freely. However, we did not observe this expected pattern for LD for the most common homozygous cluster in paracentric inversions. We found that across paracentric inversions, the most common homozygous cluster often had similar LD levels to all genotypes (SI Appendix, Fig. S3). Conversely, as pericentric inversions encompass the centromere, high LD was observed within and between clusters as expected. Overall, our results are consistent with signatures of low recombination with high LD in all predicted centromeric regions and most predicted inversion regions.

To corroborate these inversions with PacBio high fidelity long-read sequencing (HiFi), we completed pairwise comparisons between the ‘Alphonso’ reference genome used for the local PCA and haplotype sequences of three genomes where all the chromosomes were assembled telomere-to-telomere (SI Appendix, Fig. S5). These three genomes corroborated all but one of the local PCA identified inversions, providing strong evidence for the prevalence of these inversions within the *M. indica* gene pool. Interestingly, in contrast to paracentric inversions, multiple HiFi inversions were often identified within the local PCA pericentric inversions (Table 1). This indicates that centromeres are indeed hotspots for the origin and persistence of inversions, where the local PCA approach identifies multiple similar, closely located pericentric inversions as a single block of differentiated genomic structure.

Overall, chromosomal inversions and regions of genetic differentiation accumulate in the centromeric areas of *M. indica*. This observation implies that low recombination regions could facilitate the preservation of genetic variation that might otherwise be susceptible to disruption through recombination. Such hotspots of genetic differentiation might contribute to variation in key breeding traits.

### Inversions harbor loci for key breeding traits

To understand the potential role of identified inversions in shaping genetic variation for key mango breeding traits, we assessed the joint contribution of the inversions to trait variation using two approaches. The first approach fitted all inversions as fixed effects in a linear model and the second approach tested the importance of each inversion in fitting an iterative random forest model for each trait. The key mango traits included fruit blush color, fruit firmness, fruit weight, total soluble solids (an indicator of sweetness) and trunk circumference (an indicator of tree size). We found that 11 of the 17 inversions are associated with at least one of these traits (Table 1 and SI Appendix, Table S4). Both models show that the inversions jointly explain a large amount of the genetic variation in blush color (33%) and total soluble solids (21%), but a moderate amount for trunk circumference (16%) and fruit weight (12%), and very low for fruit firmness (2%) (SI Appendix, Table S5). Among these inversions, miinv6.0 shows an additive relationship with blush color and weight, where accessions in genotype cluster 0 have more blush and heavier fruit than the other clusters (Fig. 2). Selection for miinv6.0 cluster 0 accessions could be a simple approach to increase fruit blush color and weight in *M. indica*. These results suggest that inversions facilitate the maintenance and persistence of allelic combinations that contribute to variation in these key breeding traits.

**Fig. 2.**
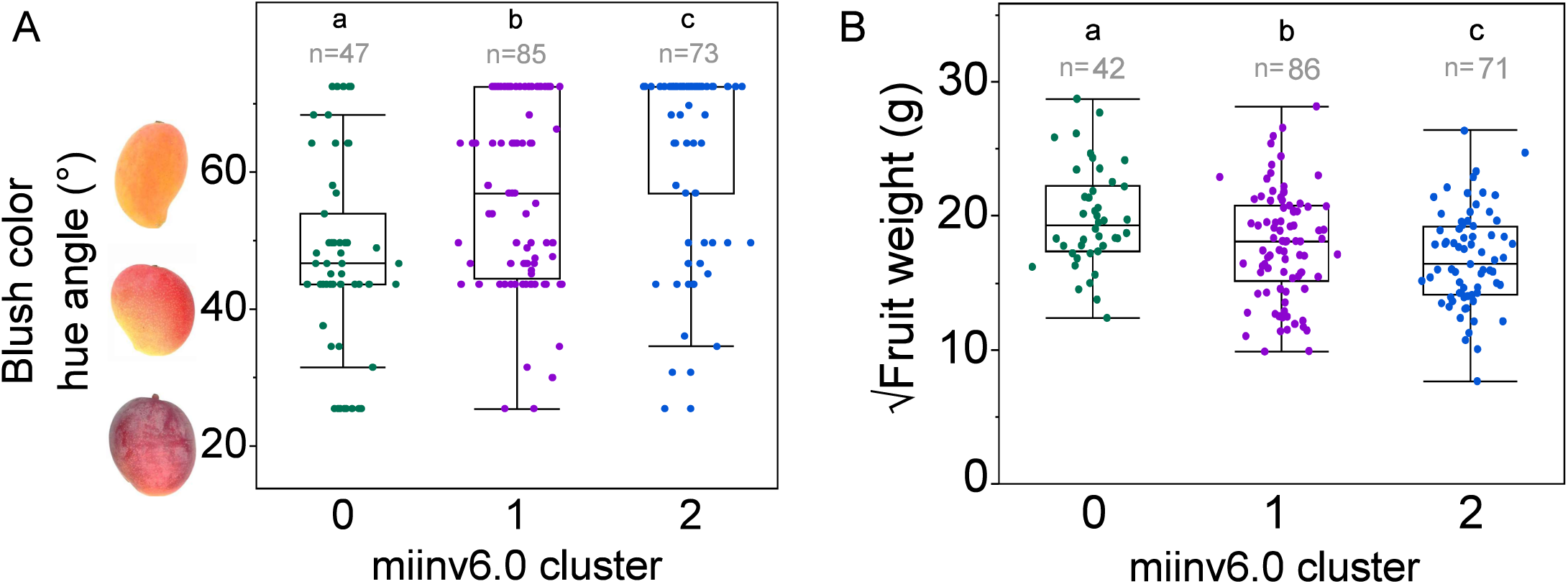
Inversions are associated with trait variation. miinv6.0 segregates with A) fruit blush color hue angle (°) and B) fruit weight (g). The square root of the fruit weight was used to normalize the data for analysis. Clusters were predicted using the k-means clustering from the local PCA. Letters denote significance level of α = 0.05. From the joint linear model, miinv6.0 has an additive effect of +6.8 hue angle (°) and -2.2 √fruit weight (g) as cluster number increases.

To identify the genes underlying these key breeding traits and their association with the inversions, we examined the distribution of loci discovered using genome-wide association studies (GWAS). The GWAS for blush color and total soluble solids showed a single major cluster of associated SNPs (Fig. 3), or quantitative trait loci (QTL). One of the most associated SNPs within the QTL for blush color (miBCSNP15:10730980) explains a very large portion of the genetic variation in blush color (45%; SI Appendix, Fig. S6) and for total soluble solids, the top SNP (miTSSSNP5:7109828) explains 19% of the genetic variation (SI Appendix, Fig. S7). On the other hand, fruit firmness, trunk circumference, and fruit weight associated SNPs were spread throughout the genome, with multiple QTLs (Fig. 3), indicative of highly polygenic traits. Overall, we found 44 (10.1%) GWAS SNPs within the identified 17 inversions, which does not exceed what you would expect by chance (quantiles 2.5% = 10.4% and 97.5% = 10.6% from 1,000 bootstraps). Even though there is no enrichment of GWAS signals across all inversions, one inversion (miinv5.0) is unusually rich in harboring GWAS SNPs (quantiles 2.5% = 0.96% and 97.5% = 1.01% from 1,000 bootstraps). miinv5.0 contains 32 (7.36%) GWAS SNPs for total soluble solids within miTSSQTL5 (Fig. 3). The absence of enrichment of GWAS SNPs within other inversions might be due to many small effect loci not detectable with a low powered GWAS.

**Fig. 3.**
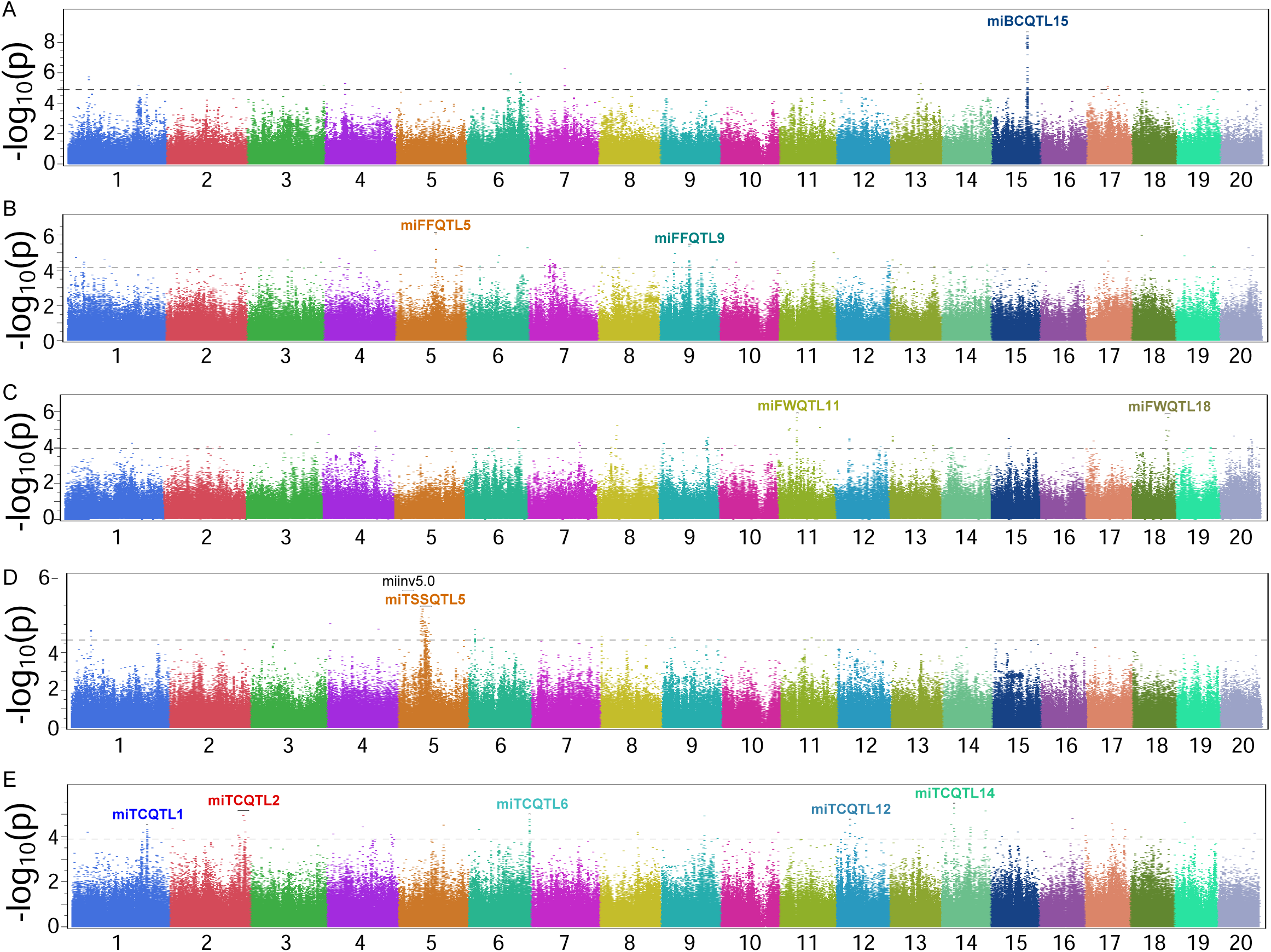
Manhattan plot of A) fruit blush color, B) fruit firmness, C) fruit weight, D) total soluble solids and E) trunk circumference. The strength of association of genomic markers with the trait (-log10(p)) across all 20 chromosomes. miinv5.0 overlaps with miTSSQTL5.

We assessed the enrichment of genes within inversions to evaluate the contribution of each inversion to function. Here, we focused on fruit blush color as the pathways involved in pigmentation are well known. Among the four inversions associated with blush color identified from the linear joint analysis (miinv1.0, 3.0, 6.0 and 9.0), three exhibit strong enrichments of relevance when using the top 100 ranked neighbor genes determined by multi-omic network analysis (see Methods; Dataset S1). Specifically, miinv9.0 is enriched for genes influencing levels of carotenoid, known to impart the orange or yellow colors on mango skin (35). Conversely, miinv1.0 is enriched for genes influencing levels of flavonoid and miinv6.0 for jasmonic acid signaling. Preharvest spraying of jasmonic acid is known to increase blush through upregulation of the flavonoid - anthocyanin (36, 37). Consistent with the major role of anthocyanin in producing red blush, the QTL in chromosome 15 from the blush color GWAS (Fig. 3), contains a transcription factor MYB114-like (MYB1) that is known to affect anthocyanin production in many cultivated species. Kanzaki*, et al.* (38) identified MYB1 as a regulator of light dependent red coloration in mango. This implies that anthocyanin regulation is key to red blush in mango and the genetic variation underlying this is spread across multiple chromosomal inversions and other pockets of the genome such as chromosome 15 (Fig. 4).

**Fig. 4.**
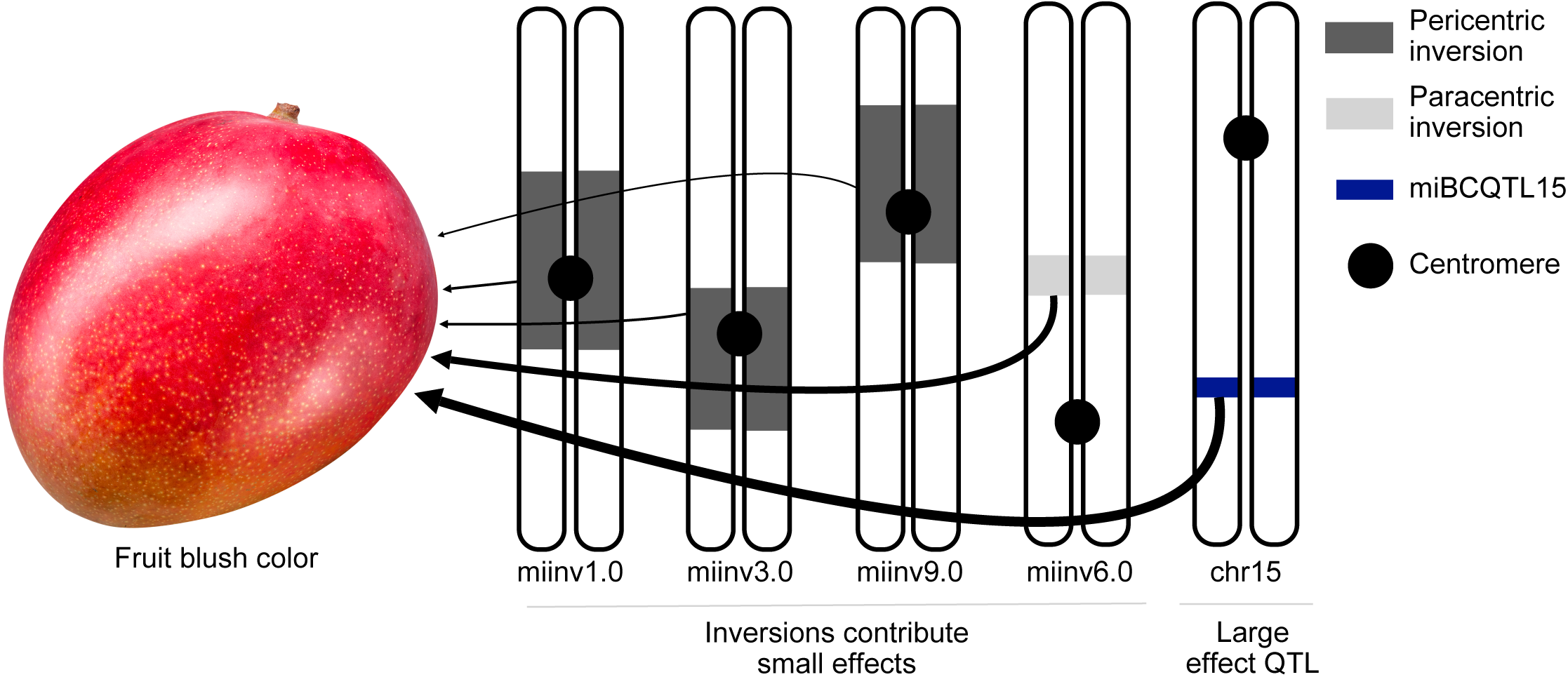
The genetic architecture of fruit blush color. Inversions, which are either pericentric (located across the centromere) or paracentric (located on one side of the chromosomal arm), contain many loci that contribute to the variation in fruit blush color. The quantitative trait loci (QTL) on chromosome 15 contains few large effect loci. The size of the arrow represents the magnitude of the contribution to fruit blush color.

These results suggest that loci of large effect together with allelic variation contained in chromosomal inversions help create specific combinations of alleles that contribute to key breeding traits in mango. As such, polygenic selection aided by chromosomal inversions has likely contributed to trait differentiation among mango accessions from origins across the world. The potential for selecting large chromosomal rearrangements, such as inversions, instead of SNPs, opens a novel and potentially efficient avenue for selection of favorable polygenic traits.

### Centromeres and large pericentric inversions have high deleterious scores

To investigate the distribution of deleterious alleles in the Australian mango gene pool that could antagonize selection for important traits, we implemented Sorting Intolerant From Tolerant (SIFT) (39) that identifies damaging mutations based on sequence homology. Here, we defined a deleterious score as 1-SIFT, where > 0.95 are putative deleterious mutations. We found that 19% (n=37,503) of genic sites across the genome are deleterious, with the remaining genic SNPs being non-damaging or tolerated (SI Appendix, Table S6). As expected, deleterious alleles are at low frequencies in the mango gene pool (SI Appendix, Fig. S8).

To test the prediction that deleterious alleles accumulate in regions of low recombination in the *M. indica* genome, we assessed the average deleterious score in centromeric and inversion regions. We found that sites within 1Mb of the predicted centromere have a higher deleterious score compared to the rest of the genome (mean deleterious score: centromere = 0.506 and non-centromere = 0.484; quantile 97.5% = 0.486 from 1,000 bootstraps). This is consistent with regions of low recombination accumulating deleterious alleles, where selection cannot purge them easily. The deleterious score of pericentric inversions was also higher relative to the rest of the genome (mean deleterious score: pericentric inv = 0.501 and non-pericentric inv = 0.485; quantile 97.5% = 0.487 from 1,000 bootstraps), but deleterious scores for paracentric inversions were not significantly different from background levels (mean deleterious score: paracentric inv = 0.483 and non-pericentric inv = 0.485; quantile 97.5% = 0.487 and 2.5% = 0.483 from 1,000 bootstraps). Specifically, deleterious scores varied considerably across the 17 inversions, where miinv4.0 had the highest deleterious score (0.597) and miinv1.1 the lowest (0.362; Table 1). For pericentric inversions, the distribution of deleterious scores is driven by inversion size, where large pericentric inversions have high deleterious scores (Fig. 5). This suggests that the size of the mutational target in non-recombining regions of the genome might contribute to the accumulation of deleterious alleles.

**Fig. 5.**
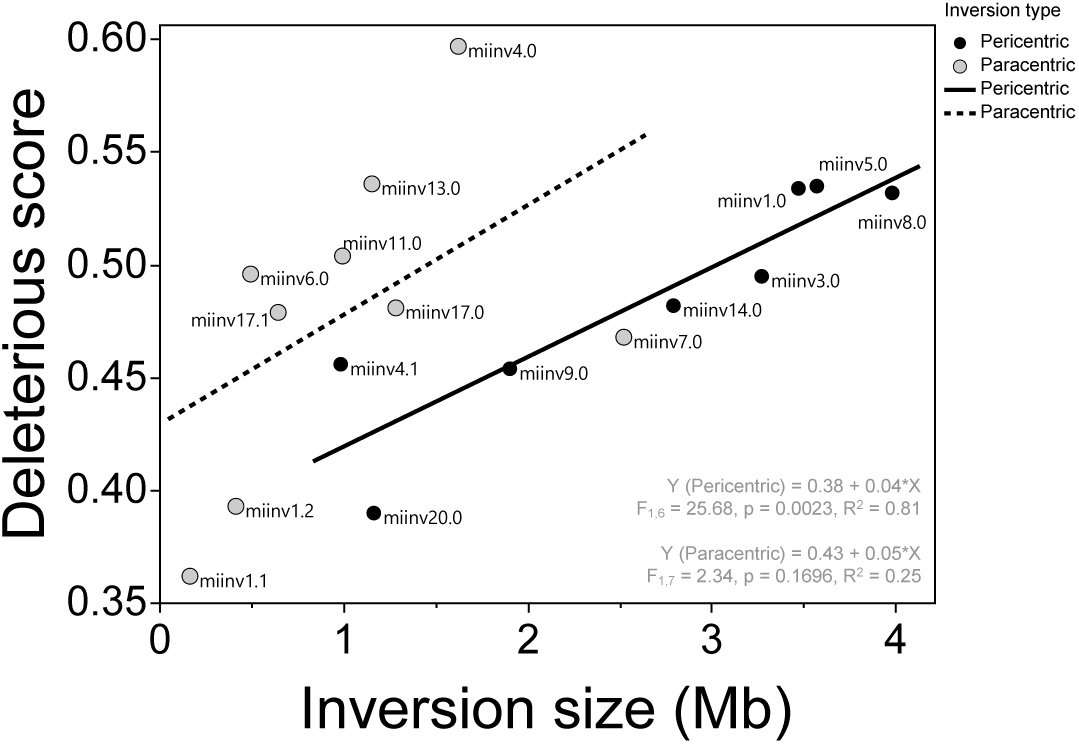
Inversion size predicts deleterious score in *M indica*. High deleterious scores (1-SIFT) reflect more deleterious alleles. The inversions are classified as paracentric (located on one side of the chromosomal arm) and pericentric (located across the centromere).

### QTL regions under selection display low deleterious scores

We have shown that harmful alleles often gather in regions with low recombination in *M. indica*. However, a critical question arises: can selection effectively overcome the drag of these deleterious alleles to increase the frequency of desirable alleles? To explore this, we focused on a large QTL for total soluble solids (miTSSQTL5 = 3.2Mb) that has a 1Mb overlap with a sizable pericentric chromosomal inversion (miinv5.0 = 3.6Mb). Despite this inversion being laden with deleterious alleles, the QTL for total soluble solids has a smaller deleterious score compared to the genome background (mean deleterious score: miinv5.0 = 0.535, miTSSQTL5 = 0.482, 1Mb overlap = 0.507 and background = 0.485; quantiles 2.5% = 0.483 and 97.5% = 0.487 from 1,000 bootstraps). This suggests that strong selection might have occurred for desirable alleles in this QTL, overcoming linkage to detrimental alleles.

To further understand the evolution of this QTL (miTSSQTL5), we evaluated the frequency of the main locus for this trait - the top GWAS SNP for total soluble solids (miTSSSNP5:7109828). This SNP lies within miinv5.0 (330kb from the end) and exhibits a significant association with the inversion (Fisher’s exact test, n = 225, P < 0.0001), where inversion cluster 2 is fixed for the most common allele at this site. Inversion cluster 0 and 1 have low frequencies of the alternate allele. The common allele (frequency in the gene-pool = 0.92) is additively associated with a favorable high total soluble solids value (40), suggesting the desirable allele is close to fixation in the gene pool. These results suggest that intense selection for high total soluble solids could be driving this allele to fixation in *M. indica*.

Consistent with signatures of selection in the mango genome, QTLs for traits that have been actively selected for in mango breeding programs, like fruit blush color, total soluble solids, and fruit weight, show a reduced deleterious score (SI Appendix, Table S7). Our findings suggest that the power of selection in breeding has not only shaped desirable traits but has incidentally minimized potentially harmful alleles in cultivated mango varieties.

## Discussion

Chromosomal inversions are known to play a large role in trait evolution in nature (1–5, 41, 42), yet inversions are rarely used as a tool for breeding (see exceptions in maize (43) and rice (44)). In many cases, selection for breeding traits has likely led to the incidental change in frequency of an inversion (12). Intentionally harnessing the influence of these low recombination regions could drive major shifts in key agronomical traits. Our study is a step forward for understanding the role of chromosomal inversions in maintaining breeding trait variation. Here, we provide empirical evidence in *M. indica* to suggest that chromosomal inversions are a major force driving breeding trait evolution, through the aggregation of small effect alleles in regions of low recombination.

Further, we find these chromosomal inversions are often located across centromeres, where large pericentric inversions are accumulating putative deleterious alleles. Although these deleterious alleles can lead to reduced fitness, our data also suggests that pockets of deleterious alleles could be potentially removed through selection for desirable traits. The results from this study suggest selection for desirable inversions could be a simple effective strategy for breeding for polygenic traits.

Pericentric inversions amplify the low recombination rates near centromeres making it unlikely to generate haplotype recombinants. This severely limits the potential for the inversion to be broken up by gene flux and could explain why polymorphic inversions were commonly located within centromere regions here in mango and other species such as deer mice (1). The severely reduced recombination can also help explain why pericentric inversions are large in mango, maize (43), and sunflower (45), and have stronger genetic signatures and divergence compared to paracentric inversions, as shown in this study. Paracentric inversions do not have the same level of suppressed recombination, so there are more opportunities for recombinants to arise within the inversion via gene flux. The centromere-amplifying effect might provide pericentric inversions the ability to preserve allelic combinations as stable polymorphisms, while paracentric inversions might be more transient in the polymorphic state.

The severe reduction in recombination in these large pericentric inversions might also contribute to the accumulation of more deleterious alleles. Our results are consistent with sorghum and maize, where predicted deleterious mutations were frequent in severely suppressed recombination regions, such as centromeres (20, 23). However, in regions of the genome that are not as severely suppressed such as paracentric inversions, we found no overall accumulation of deleterious alleles within inversions, which is similar to deer mice (1) and sunflower (45) that also used the local PCA approach to identify polymorphic inversions. Harringmeyer and Hoekstra (1) suggest inversion homozygotes allow recombination to be uninterrupted, thus implying that the inverted regions will gradually purge deleterious alleles as inversion haplotypes increase in frequency in the population. Phylogenetic analysis of the inversions with wild and domesticated *Mangifera* will help elucidate the age of these inversions and whether they facilitated the accumulation of favorable or deleterious loci.

Natural and artificial selection can purge deleterious alleles from regions of the genome, but it can also drive the accumulation of deleterious alleles. In populations with substantial haplotype diversity, selection can replace the harmful haplotypes with the favorable haplotypes (22, 46). On the other hand, deleterious alleles genetically linked to loci targeted by selection can spread in the population (21, 28, 47). In the absence of recombination, these linked regions span larger distances which increases the probability of deleterious alleles evolving together with selected alleles. Our study found that QTLs for fruit blush color, weight and total soluble solids have reduced deleterious scores. This result mirror those by Zhu et al. (46) that found strong artificial selection was correlated with low genetic load in rice, soybean, tomato, grape, and pineapple, with the exception of centromeric regions. These results highlight the amplifying effects of centromeres on reducing recombination rate, which is particularly evident in mango in large pericentric inversions that harbor both deleterious and favorable loci (e.g. miinv5.0, miinv1.0 and miinv3.0). After close examination of the one QTL overlapping an inversion, we see that selection for the total soluble solids QTL likely helped remove deleterious loci in tight linkage. These results suggest that selection for loci within QTL peaks may induce linked purifying selection against nearby deleterious variants, even in regions of low recombination. Overall, examining genome structure dynamics surrounding sites under selection provides unique insights into the tensions between maintaining desirable variation and removing deleterious mutations.

The clustering of loci for key breeding traits within inversions implicates a broader role of structural mutations like inversions in orchestrating complex, polygenic trait architectures. Targeting of inversions containing adaptive loci for polygenic traits has been observed in nature (41). For breeding, tracking chromosomal rearrangements may offer a more efficient approach than using individual SNPs as we can leverage the combination of alleles held together by low recombination. For example, we suggest a simple approach to increase mango fruit blush color and weight by selecting for individuals identified in cluster 0 of the inversion on chromosome 6. This can be combined with selection for other inversions to further increase trait values. To identify these inversions in untested accessions, inexpensive sequencing methods such as targeted capture and low coverage sequencing can be completed to identify some of the SNPs in the region. Future studies should analyze the contribution of these inversions to trait variation in different breeding programs to understand the contribution of the environment and management practices, and therefore its utility outside of the Australian mango breeding program.

Exploring the interconnections between centromeric suppression of recombination, chromosomal inversion dynamics, maintenance of breeding trait variation, and accumulation of deleterious mutations provides fundamental insights into genome architecture evolution. These findings deliver direct applied outcomes surrounding optimal tracking of chromosomal inversions to select for complex mango breeding traits rather than individual loci. Our research also carries broader relevance for the ubiquity of centromeric expansive effects in shaping chromosome-wide variability and trade-offs between positive and negative selection over generational timescales.

Overall, our results can facilitate the elimination of deleterious effects and enable the selection of favorable traits, resulting in remarkable improvements in mango varieties. Moreover, the insights garnered from our research hold the potential for application to other tree crop species, thereby enhancing overall crop productivity.

## Materials and Methods

### Plant material

We selected 225 *Mangifera indica* from the gene pool collection of the Australian Mango Breeding Program managed by the Department of Agriculture and Fisheries at Walkamin Research Station, Queensland (17.1341°S, 145.4271°E). These samples were originally imported from 24 countries across five geographical regions and grafted onto the uniform polyembryonic rootstock, Kensington Pride. The samples with one parent known are from an open-pollination cross (Dataset S2).

### Genotyping

We extracted DNA from 225 mango accessions collected from Walkamin Research Station. DNA was extracted from young leaves following Healey, Furtado, Cooper and Henry (48) CTAB (cetyltrimethylammonium bromide) method with the following modifications. 5 g of young leaves were ground using a mortar and pestle. 16 mL nuclear lysis buffer and 4 mL Sarkosyl solution were added and gently mixed with the leaf tissue. Tubes were incubated for 2 h at 65°C in a water bath with periodic mixing. The RNAse digest (step 5) was moved to day 2 and allowed to incubate at room temperature for 1 h. 5 M NaCl was added to the final concentration of 0.25 M and mixed well. 0.35 vol 100% EtOH was added and quickly mixed. The samples were incubated on ice for 10 mins and centrifuged for 15 mins at 10°C at 9,000 rpm. The solution was transferred to a new tube and an equal volume of chloroform was added and gently inverted 50 times.

Samples were again centrifuged for 15 mins at 10°C at 9,000 rpm. The upper phase of the solution was transferred to a new tube. One volume of isopropanol was added and the tube was inverted to mix and centrifuged for 15 mins at 10°C at 13,000 rpm. The solution was removed and 500 μl 70% EtOH was added and centrifuged for 15 mins at 10°C at 13,000 rpm. The solution was removed, and the dry pellet was resuspended in TE buffer. DNA quality and quantity was assessed using qubit, nanodrop, and a 1% agarose gel. For the nanodrop 260/280, samples were between 1.7 – 2 and 260/230 > 1.4.

We performed whole genome sequencing on 225 mango samples at the Ramaciotti Centre for Genomics, UNSW, New South Wales, Australia. Sequencing libraries were generated using the Illumina DNA-prep kit and subjected to 150bp PE sequencing on one S4 flow cell of a NovaSeq 6000 Illumina sequencer. Sequencing was undertaken to obtain a minimum data depth for each of the samples; 41 mango samples with an expected coverage of 40X and 184 mango samples with an expected coverage of 15X. The size of the raw data provided by the Ramaciotti was more than requested.

### Alignments and variant calling

To obtain SNPs for downstream analyses we joint-called SNPs using the GATK4 software package and best practices developed by the Broad Institute for variant detection. The publicly available GATK pipeline (https://gencore.bio.nyu.edu/variant-calling-pipeline-gatk4/) was heavily modified to include functionality for read processing, deduplication, quality control, parallel processing and joint-calling. Trimmed paired-end reads and singletons from 225 re-sequenced *M. indica* genomes (25x average coverage, 100x maximum coverage) were aligned to the *M. indica* cv ‘Alphonso’ reference genome (NCBI GenBank assembly accession: GCA_011075055.1) (49) using BWA MEM v0.7.17 (50) with ‘-v 3 -Y -K 100000000 -M’ parameters. This approach produced 44,125,383 SNPs (1 every 9 bases).

### Quality filtering

The data was quality filtered using the following parameters in VCFtools v0.1.17 (51): minimum genotype quality of 20; minimum depth per sample of 5; missing data per site 50% (initial relaxed threshold); maximum mean depth of 50 for all sites (removal of paralogues); and a missing data per site 20% (stringent threshold). All accessions have more than 85% of SNPs. We then used GATK VariantFiltration for standard quality filtering which removed: QualByDepth < 2; quality score < 30; StrandOddsRatio > 3; FisherStrand > 60; RMSMappingQuality < 40; MappingQualityRankSumTest < -12.5; ReadPosRankSum < -8. Indels were then removed using VCFtools v0.1.17(51) and contigs that did not align to a chromosome were removed using BCFtools v1.12 (52).

### Predicting centromere locations

To predict centromere locations on each chromosome, we used RepeatOBserver (33). Centromeres were identified as the regions on each chromosome where the most repeat lengths reached a minimum abundance, implying the dominance of a single centromeric repeating sequence in this window. Specifically, chromosomes were divided into 2Mb windows. The total abundance of each repeat length was determined in each window. The window where each repeat length minimized its abundance was identified. A histogram of the number of repeats minimizing in any given window was plotted (SI Appendix, Fig. S2). The peak in the histograms is the predicted centromere location.

### Local principal component analysis

To prepare the data for local PCA, we removed invariant sites using GATK SelectVariant v4.2.5 (53). We removed sites with greater than 95% missing data using VCFtools v0.1.17 (51). We converted the SNP dataset to BCF format with BCFtools v1.12 (52) and 15,957,988 SNPs were used for analysis.

To detect genomic regions of unusual population structure we performed a local PCA using the R package ‘lostruct’ v0.0.0.9000 (18). We performed a PCA on windows of 1,000 SNPs across the genome using the lostruct function eigen_windows. To measure similarity between these PCA matrices (for the first two PC values) we applied the pc_dist function in lostruct. We extracted the first 40 Multidimensional Scaling (MDS) coordinates in R v4.1.3. To visualize clustered outliers with extreme population structure relative to the genome wide average we plotted the MDS coordinates along each chromosome. Following Huang, Andrew, Owens, Ostevik and Rieseberg (3), we defined outliers as windows with MDS values greater than 4 standard deviations from the genome wide mean. We combined outlier windows with less than 10 windows between them to form clusters, with clusters requiring a minimum of four outlier windows to be considered. To determine if the distribution of outliers deviated from random expectations, we performed 1,000 permutations of windows with a significance value of p < 0.01. For each MDS coordinate we analyzed outlier clusters in both positive and negative directions separately. For chromosomes with outlier clusters on multiple MDS coordinates we collapsed those with a Pearson’s product moment correlation coefficient greater than 0.8 and selected the MDS coordinate with the larger number of outliers. We defined the coordinates of each inversion candidate by the start of the first outlier window in the cluster and the end of the last outlier window.

To find outlier clusters for which samples were divided into three groups (representing two homozygous and a heterozygote genotype, as expected for a polymorphic inversion) we performed PCA with the R function ‘kmeans’. The maximum, minimum and middle of the PC1 scores were used as the centers for each of the three clusters and the discreteness of the clusters was measured as the proportion of the between-cluster sum of squares over the total sum of squares.

### Homozygosity

To identify signatures of chromosomal inversions from the 28 outlier windows identified in the local PCA, we calculated observed homozygosity within each outlier window using the --het flag in VCFtools v0.1.17 (51). Outlier windows with local PCA clusters 0 and 2 (predicted homozygous clusters) that are significantly higher in homozygosity compared to cluster 1 (predicted heterozygous cluster) were considered as potential chromosomal inversions in this study.

### Genetic differentiation

We measured genetic differentiation (FST) between local PCA clusters 0 and 2, the predicted homozygous clusters. We removed SNPs with MAF < 0.01 and applied --weir-fst-pop for 10kb windows in VCFtools v0.1.17 (51). The weighted FST output was used for all analyses.

### Nucleotide diversity

Nucleotide diversity (π) was estimated on all 263 million sites (including invariant) across 10kb non-overlapping windows using pixy v1.2.7.beta1 (54) for each inversion cluster. Pixy was used due to its unbiased estimates of nucleotide diversity in the presence of missing data.

### Linkage disequilibrium

We calculated mean linkage disequilibrium (LD) across the genome with pairwise r2 for all SNPs in non-overlapping 200kb windows using plink v1.9 (55).

### PacBio high fidelity long-read sequencing

To verify the chromosomal inversions identified using the local PCA method, we completed genome comparisons between *M. indica* cv. ‘Alphonso’ (CATAS_Mindica_2.1) and three telomere-to-telomere genomes created using PacBio high fidelity long-read sequencing.

Methodology for DNA extraction, library preparation and genome creation can be found in Wijesundara et al. (56). Genome comparison between *M. indica* cv. ‘Alphonso’ (CATAS_Mindica_2.1) and Irwin, Kensington Pride and *M. laurina* haplotypes were conducted separately to identify inversions between genomes. The genomes were aligned using MUMer v4 (57) with parameters maxmatch -c 100 -b 500 -l 50. Each alignment was filtered using the delta-filter option implemented in MUMer with parameters -m -i 90 -l 100. Sequence differences analysis was performed using the Synteny and Rearrangement Identifier (58) and visualized using plotsr v1.1.0 (59).

### Phenotyping

We phenotyped five key breeding traits in the Australian mango gene pool – fruit blush color, fruit firmness, fruit weight, total soluble solids and trunk circumference (Dataset S2). All fruit samples were harvested from trees after maturity (> five years old). Each accession had ten fruits assessed from the outside of each tree, where the fruit is exposed to full sun. After harvest the fruit was de-sapped, where they were dipped for 5 minutes in a hot fungicide dip at 52°C with 1.0 ml L-1 Fludioxonil (230g/L). Fruit blush color was assessed in 205 accessions annually for at least 2 years by categorizing the fruit blush color as either: no blush, orange, pink, red or burgundy. The average hue angle (°) for each blush color category was calculated using a Konica Minolta spectrophotometer (colormeter) in the LAB color space (L*= lightness; a* = redness; b* = yellowness) and the formula: h = arctan(b*⁄(a*)). Fruit firmness was assessed in ripe fruit of 199 accessions using an analogue firmness meter. The fruit was placed into a V-shaped metal stand and a load of 50 g placed on the fruit for 30 s. The vertical displacement (mm) of the fruit over this time was recorded and the mean from ten fruit was used. Fruit weight (g) in 199 accessions was assessed in ten mature fruit prior to ripening. The fruit was weighed on a digital laboratory balance and the mean weight of the ten fruit was used. Total soluble solids was assessed using the juice from one cheek for 198 accessions. A cheek was placed on a digital refractometer and total soluble solids was measured on the BRIX scale. The BRIX values from ten fruit were averaged for each accession. Trunk circumference (cm) was assessed on 178 accessions by measuring the scion 10 cm above the graft on trees that are 12 years old. Trunk circumference and fruit firmness are normally distributed, while total soluble solids underwent log10 transformation and fruit weight square root transformation. Blush color was not normally distributed and unable to be transformed.

### Joint contribution of inversions on trait value

We considered each of the 17 putative inversions as genetic markers and genotyped them as 0, 1 and 2 (homozygous, heterozygous, and homozygous alternate states according to the local PCA clustering described above) in 225 mango accessions. We then fit a linear model for each of the breeding traits using the 17 inversion markers as fixed effects. Significant inversions were recorded for each trait, and the total variance explained by all inversions for a trait was given by the adjusted R^2^ value of the model. We also fit a non-linear iterative Random Forest model for each trait using ‘iRF’ v3.0.0 in R (60) and selected the top 3 most predictive inversions for each trait according to sorted variable importance.

### Genome wide association study

To identify the putative loci underlying three key mango traits, we performed a genome wide association study (GWAS) in plink v1.9 (55). Rare alleles (MAF < 0.01) and non-independent SNPs (window size of 50kb, step size of 10 and r^2^ threshold of 0.1) were removed from the dataset, leaving 871,869 SNPs. To account for population structure, we performed a PCA in plink v1.9 (55) on the subset of individuals with phenotypic data - blush color (n=205) and fruit firmness (n=199), fruit weight (n=199), trunk circumference (n=178) and total soluble solids (n=198). The number of principal components included as covariates in the linear model was determined using quantile-quantile plots, where blush color = PC1-PC5, fruit firmness = PC1-PC4, fruit weight = PC1-PC5, trunk circumference = PC1-PC6, and total soluble solids = PC1-PC4. A linear association was conducted in plink v1.9 (55) for each trait independently. For blush color, we corroborated our results with a binary GWAS (no blush vs blush), which had a genomic inflation factor of 1.09 and produced similar results.

### Enrichment of GWAS loci within inversions

To determine whether the GWAS SNPs were enriched within the 17 identified inversions, we performed 1000 bootstraps in JMP v16.0.0. Of the 871,869 SNPs in the GWAS, the top 0.01% of associated SNPs for each trait was used as the random seed (n=435). Values above the 97.5% quantile were considered enriched for GWAS SNPs. We then tested for enrichment of GWAS signals in miinv5.0 using the same method.

### Functional enrichment and multi-omic context of genes within inversions

To understand the functional contribution of each inversion, we found the most functionally related genes in a multi-omic network context. First, genes within each inversion region were extracted and MangoBase (61) was used to map them to *Arabidopsis* gene ortholog identifiers. We analyzed each inversion’s *Arabidopsis* gene list for enrichment of functional annotation terms (GO BP and KEGG) using the R package ‘gprofiler2’ v0.2.2. We used RWRtoolkit v0.1 in R (62) together with an 8-layer multiplex *Arabidopsis* network containing publicly available gene-gene edges describing gene regulatory relationships, protein-protein interactions, metabolic pathways and gene knockout-effect similarity. For each inversion, we ran the *RWR_LOE* function on the multiplex network using the inversion’s gene list as seeds. This function uses the RWR network propagation algorithm to explore the multiplex topology from all seed genes jointly to score all other genes in the network according to the strength of their connectivity to the seeds. We added the top 100 highest scoring genes to the gene list of each inversion and then performed enrichment analysis on the expanded gene list.

### Identification of putative deleterious alleles

To identify putative deleterious alleles, we used Sorting Intolerant From Tolerant For Genomes (SIFT4G) (31, 32). SIFT uses sequence homology to predict whether an amino acid substitution affects protein function. To create a reference protein set, we used uniref90 from uniprot (version last modified on 2022-05-25). Genome and annotation files (GTF) for the ‘Alphonso’ (CATAS_Mindica_2.1) genome were downloaded from the National Center for Biotechnology Information (NCBI) Reference Sequence (RefSeq). To create a mango SIFT4G database, the SIFT4G_Create_Genomic_DB (https://github.com/pauline-ng/SIFT4G_Create_Genomic_DB) pipeline was followed. SIFT predictions (Dataset S3) were then created using SIFT4G (https://github.com/rvaser/sift4g) with the mango database and the VCF file containing 225 mango accessions. We defined a deleterious score as 1-SIFT, where > 0.95 are putative deleterious mutations.

To reduce the effects of reference bias we used the ancestral allele instead of the reference allele to predict deleterious mutations. We used SNPs where *Mangifera odorata* is homozygous for the reference allele to classify the ancestral state. A deleterious score is then assigned to the derived allele. We understand *M. odorata* will not directly capture the most common recent ancestor of *M. indica* as *M. odorata* has likely accumulated genetic differences since splitting from *M. indica*.

However, without using *M. odorata* to classify the ancestral state, accessions closely related to Alphonso would be identified as having fewer putative deleterious mutations as SIFT uses the difference between the reference allele and the alternate allele to determine the amino acid change that is occurring at that site. We note that we might have introduced other biases in relation to divergence from this outgroup species.

### Allelic frequency

To plot the distribution of allelic frequencies across deleterious loci, we output allelic frequencies for all SNPs with estimated deleterious scores using the --freq and --position flags in VCFtools v0.1.17 (51).

### Deleterious score statistics

To determine whether centromeres, inversions or QTL’s had a deleterious score that was significantly different from the genome average, we performed 1000 bootstraps in JMP v16.0.0 for each comparison. The random seed was based on the number of data points across inversions, centromeres or QTL’s. Values below the 2.5% quantile were considered significantly lower than the genome wide average and values above the 97.5% quantile were considered higher than the genome wide average. Deleterious scores were split into three categories based on their distance from the genome average.

To identify whether there was an association between miinv5.0 and the most associated SNP in the total soluble solids QTL, we implemented a fisher’s exact test between cluster number of miinv5.0 and the genotypes of miTSSSNP5:7109828 using JMP v16.0.0. The genotypes for this site were obtained using BCFtools query v1.12 (52) with --regions flag.

## Acknowledgments

This research was undertaken as part of the National Tree Genomics Program – Phenotype Prediction project (AS17000) which is funded by the Hort Frontiers Advanced Production Systems fund, part of the Hort Frontiers strategic partnership initiative developed by Hort Innovation, with co-investment from The University of Queensland and Queensland Government, and contributions from the Australian Government. The Australian Research Council Centre of Excellence for Plant Success in Nature and Agriculture (CE200100015) partially funded M.J.W Postdoctoral Fellowship during the execution of this work.

