## Supplementary Figures and Tables for "Centromeres are Hotspots for Chromosomal Inversions and Breeding Traits in Mango"

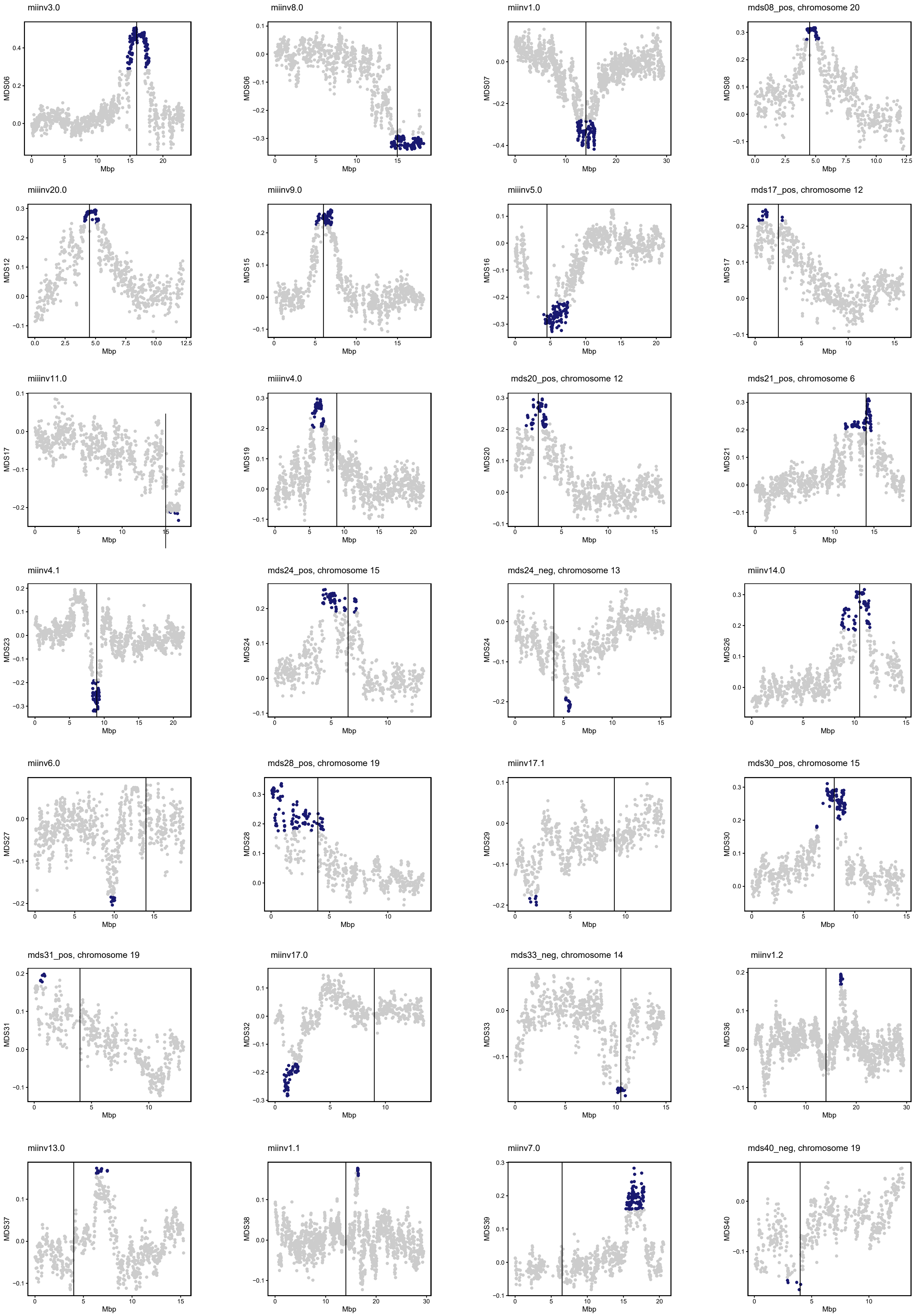


**Fig. S1.** Local PCA MDS plots. The dark blue dots show the outlier region and the black line is the predicted centromere.

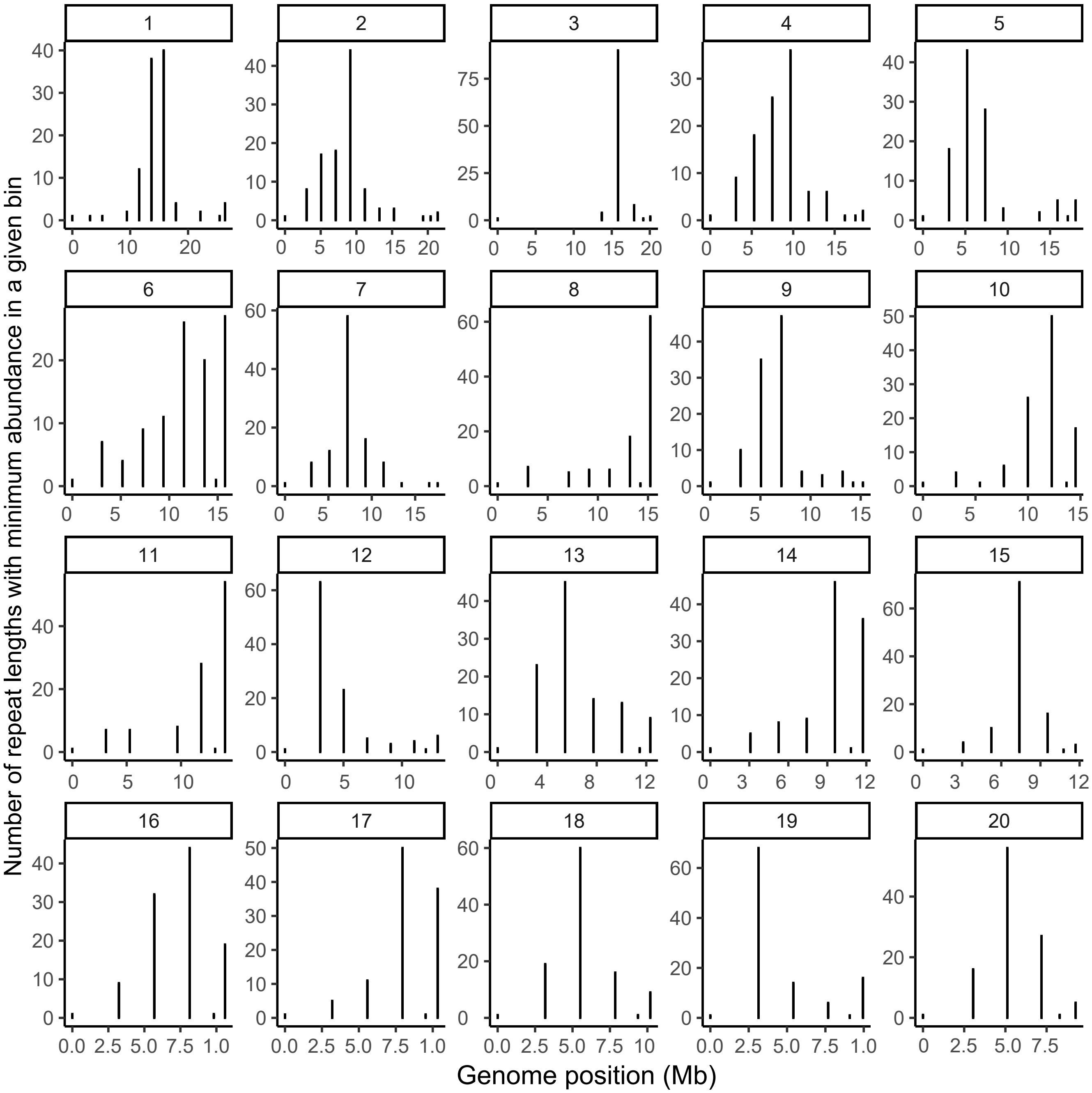


**Fig. S2.** Predicted centromere positions based on where the most repeat lengths reach minimum abundance for the 20 chromosomes in the *M. indica* cv. 'Alphonso' genome.

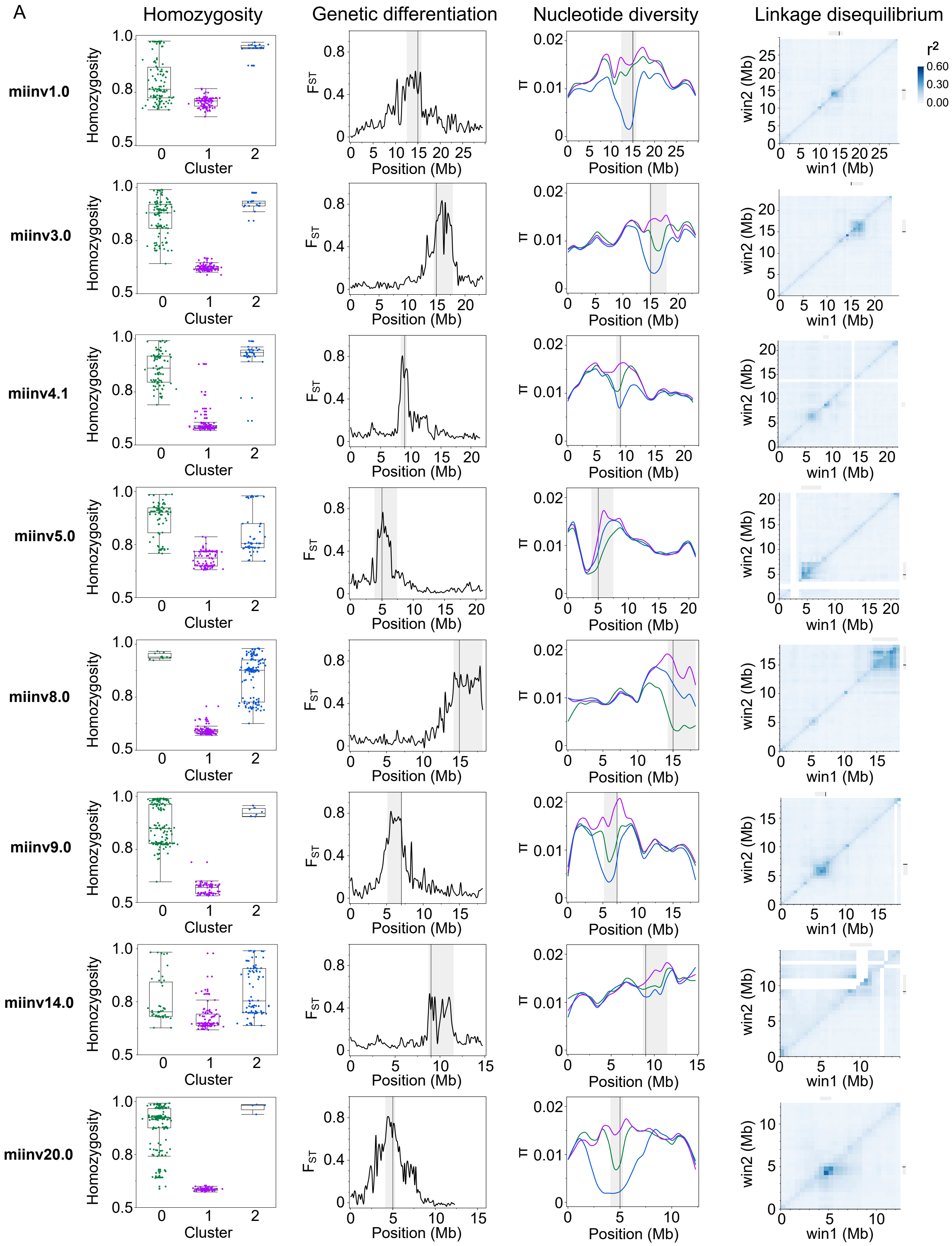
A
Homozygosity
Genetic differentiation
Nucleotide diversity
Linkage disequilibrium
miinv1.0
miinv3.0
miinv4.1
miinv5.0
miinv8.0
miinv9.0
miinv14.0
miinv20.0
Homozygosity
Cluster
$F_{ST}$
Position (Mb)
π
win2 (Mb)
win1 (Mb)
$r^2$
0.60
0.30
0.00

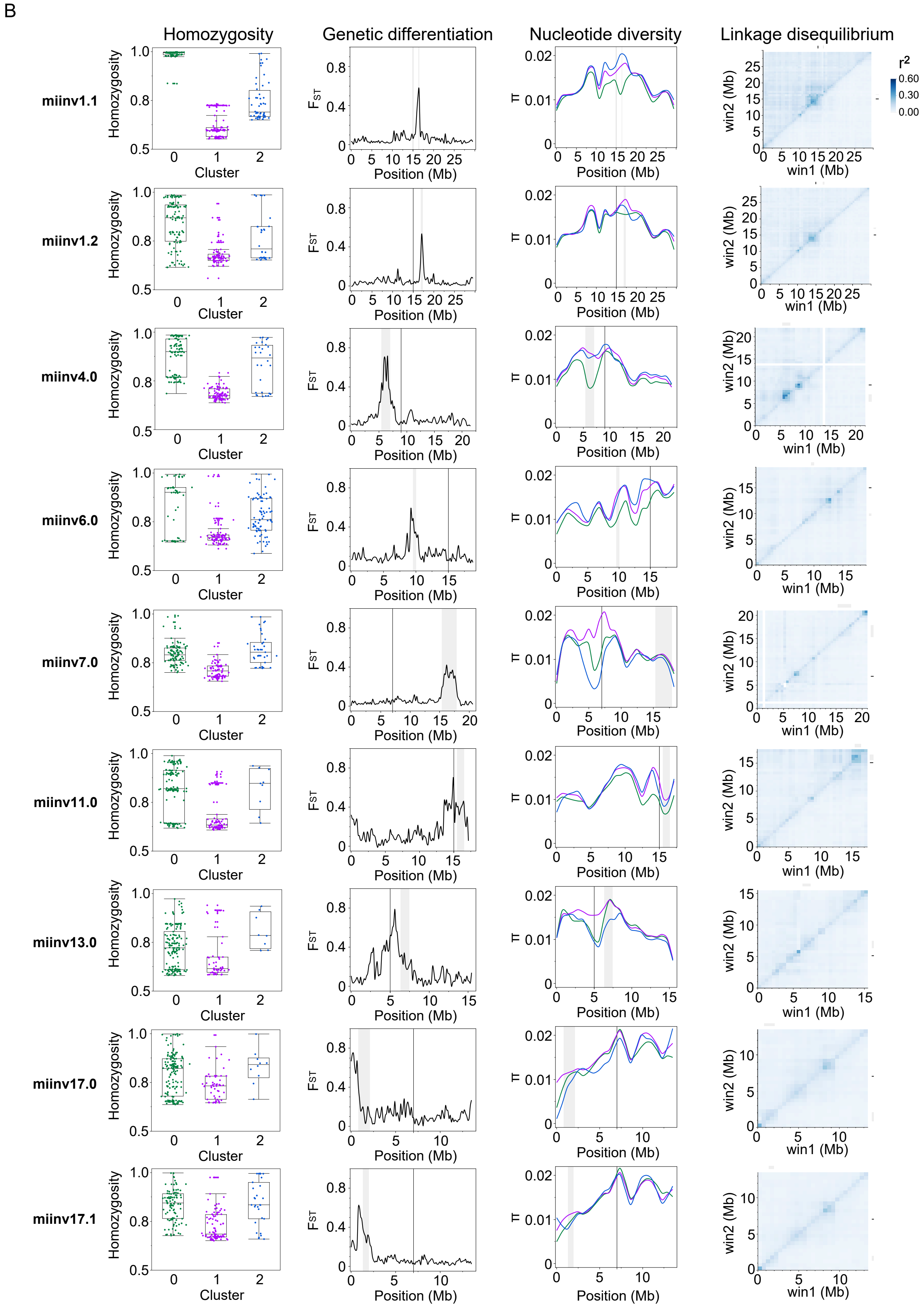


**Fig. S3**. Population genetics of A) pericentric and B) paracentric putative inversions. C) The centromere is unknown and therefore the inversion could not be classified. Homozygosity of the outlier region for the three clusters. Genetic differentiation ($F_{ST}$) between homozygous clusters, 0 and 2 across the chromosome. E) Nucleotide diversity ($\pi$) for the three clusters. F) Linkage disequilibrium heatmap of the most common homozygous group on the top triangle and all individuals on the bottom triangle (including the middle line). The black line is the predicted centromere, and the grey bar is the predicted inversion region. The colors represent the k-mean clustering (cluster 0 = green, cluster 1 = purple, and cluster 2 = blue).

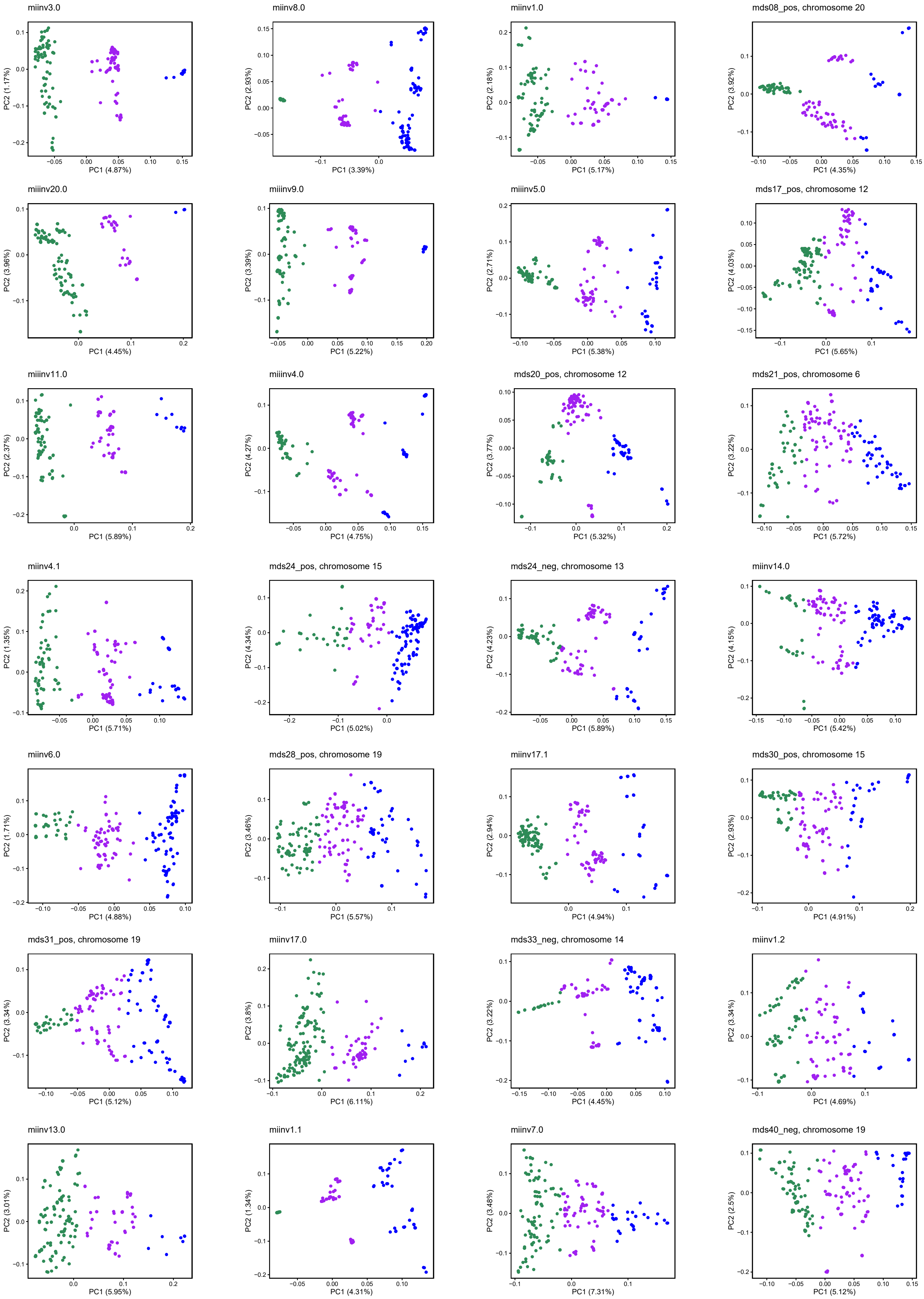


**Fig. S4**. PCA of the outlier region found from the Local PCA MDS (Fig. S1). The colors represent the k-mean clustering (cluster 0 = green, cluster 1 = purple, and cluster 2 = blue).

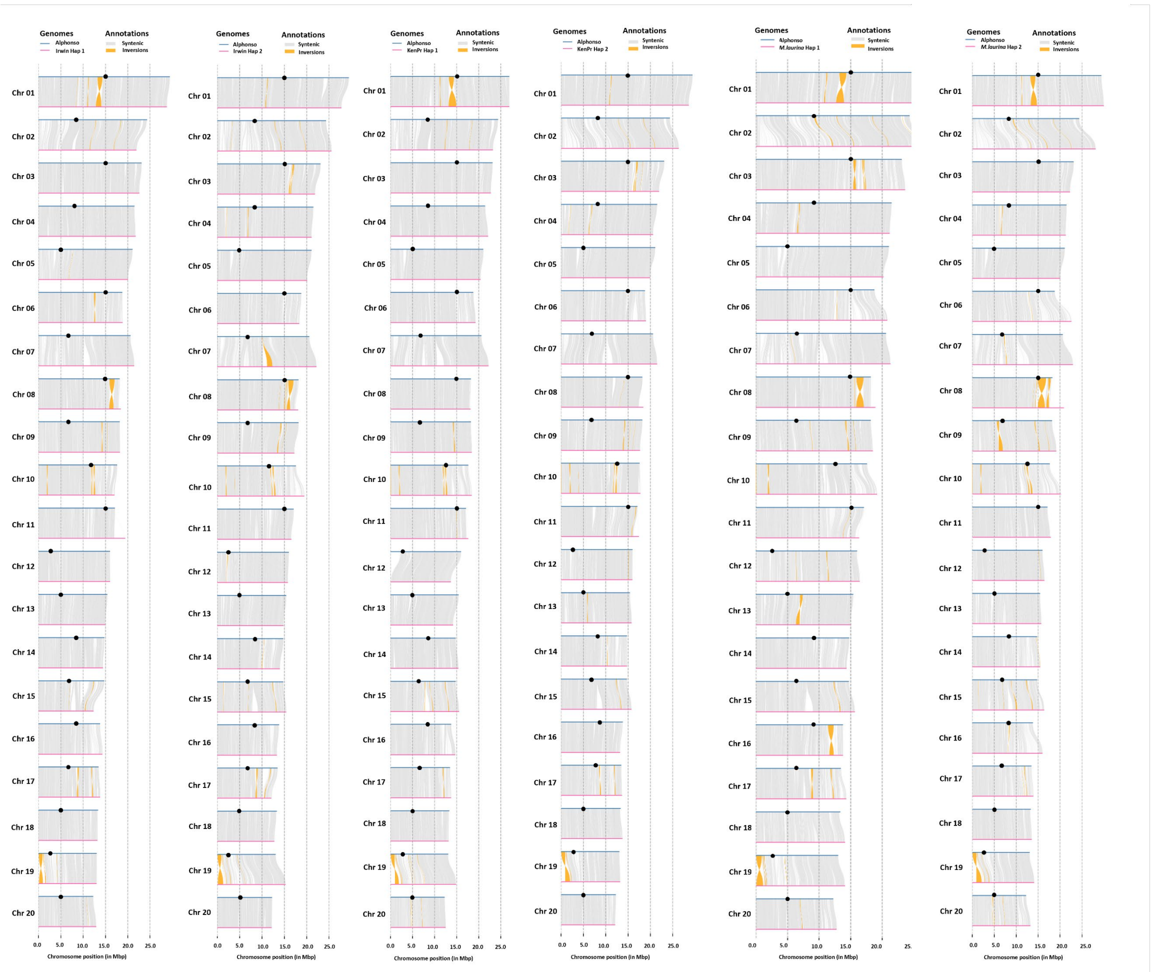


**Fig. S5.** HiFi chromosomal inversions identified in pairwise comparisons between Alphonso and the haplotypes of two *M. indica* cultivars (Irwin and Kensington Pride) and *M. laurina*, a) Irwin hap1, (b) Irwin hap2, c) Kensington Pride hap1, (d) Kensington Pride hap2, e) *M. laurina* hap1 and f) *M. laurina* hap2.

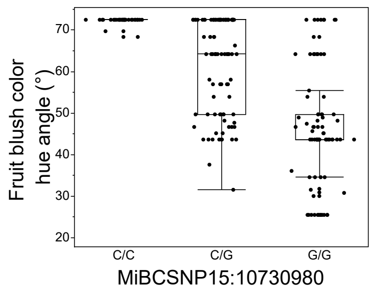


**Fig. S6.** Fruit blush color hue angle (°) for the miBCSNP15:10730980 genotypes in 190 *M. indica*. The SNP is located in the MiMYB1 gene and was significantly associated with blush color ($F_{2,187} = 77.47$, $p < 0.0001$, $R^2 = 0.45$). A high hue angle reflects a low blush color.

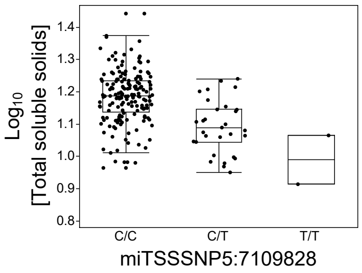


**Fig. S7**. Total soluble solids values for the miTSSSNP5:7109828 genotypes in 198 *M. indica*. Total soluble solids values were $log_{10}$ transformed to normalize the data. The SNP was significantly associated with total soluble solids values ($F_{2,195}$ = 22.95, $p < 0.0001$, $R^2$ = 0.19).

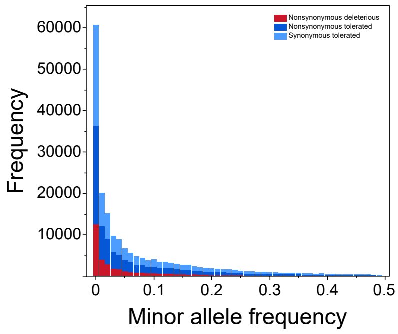


**Fig. S8**. Derived allele site frequency spectrum for nonsynonymous deleterious, nonsynonymous tolerated and synonymous tolerated for all 225 mango cultivars across variant sites only. Minor allele frequency bins of 0.01 were used.

**Table S1.** The local PCA number of outliers and discreteness of the clusters (between sum of squares (SS)) for all 28 sites identified as outlier regions.

| MDS coord. | Chr | Start | End | Region size (Mb) | N outliers | Between SS | Inv |
|---|---|---|---|---|---|---|---|
| mds07_neg | 1 | 12295528 | 15764961 | 3.47 | 115 | 0.954 | miinv1.0 |
| mds38_pos | 1 | 16336453 | 16496549 | 0.16 | 13 | 0.956 | miinv1.1 |
| mds36_pos | 1 | 16897018 | 17311598 | 0.41 | 15 | 0.868 | miinv1.2 |
| mds06_pos | 3 | 14606714 | 17874417 | 3.27 | 132 | 0.985 | miinv3.0 |
| mds19_pos | 4 | 5406761 | 7031018 | 1.62 | 64 | 0.939 | miinv4.0 |
| mds23_neg | 4 | 8327674 | 9305600 | 0.98 | 55 | 0.962 | miinv4.1 |
| mds16_neg | 5 | 3867103 | 7440236 | 3.57 | 113 | 0.955 | miinv5.0 |
| mds27_neg | 6 | 9561029 | 10053279 | 0.49 | 9 | 0.959 | miinv6.0 |
| mds21_pos | 6 | 11374502 | 14625694 | 3.25 | 77 | 0.874 | |
| mds39_pos | 7 | 15351806 | 17868042 | 2.52 | 84 | 0.883 | miinv7.0 |
| mds06_neg | 8 | 14263383 | 18239640 | 3.98 | 142 | 0.962 | miinv8.0 |
| mds15_pos | 9 | 5121940 | 7024118 | 1.90 | 72 | 0.978 | miinv9.0 |
| mds17_neg | 11 | 15503432 | 16492370 | 0.99 | 6 | 0.958 | miinv11.0 |
| mds17_pos | 12 | 379505 | 2936605 | 2.56 | 18 | 0.831 | |
| mds20_pos | 12 | 1193179 | 3368068 | 2.17 | 60 | 0.878 | |
| mds24_neg | 13 | 5216148 | 5710341 | 0.49 | 19 | 0.906 | |
| mds37_pos | 13 | 6321903 | 7467493 | 1.15 | 12 | 0.847 | miinv13.0 |
| mds26_pos | 14 | 8718458 | 11508882 | 2.79 | 78 | 0.888 | miinv14.0 |
| mds33_neg | 14 | 10160271 | 10962895 | 0.80 | 17 | 0.893 | |
| mds30_pos | 15 | 6288712 | 9084483 | 2.80 | 88 | 0.834 | |
| mds32_neg | 17 | 824929 | 2102581 | 1.28 | 47 | 0.879 | miinv17.0 |
| mds29_neg | 17 | 1308108 | 1946519 | 0.64 | 6 | 0.933 | miinv17.1 |
| mds24_pos | 18 | 4250783 | 7228351 | 2.98 | 55 | 0.865 | |
| mds28_pos | 19 | 51742 | 4485521 | 4.43 | 97 | 0.876 | |
| mds31_pos | 19 | 541569 | 928506 | 0.39 | 7 | 0.907 | |
| mds40_neg | 19 | 2879258 | 4038014 | 1.16 | 5 | 0.882 | |
| mds12_pos | 20 | 4097289 | 5258401 | 1.16 | 60 | 0.852 | miinv20.0 |
| mds08_pos | 20 | 4239755 | 5258401 | 1.02 | 50 | 0.916 | |

**Table S2.** The predicted centromere location for each chromosome in the *M. indica* cv. 'Alphonso' genome.

| Chromosome | Estimated centromere (Mb) |
|---|---|
| 1 | 15 |
| 2 | 9 |
| 3 | 15 |
| 4 | 9 |
| 5 | 5 |
| 6 | 15 |
| 7 | 7 |
| 8 | 15 |
| 9 | 7 |
| 10 | 13 |
| 11 | 15 |
| 12 | 3 |
| 13 | 5 |
| 14 | 9 |
| 15 | 7 |
| 16 | 9 |
| 17 | 7 |
| 18 | 5 |
| 19 | 3 |
| 20 | 5 |

**Table S3.** The local PCA number of outliers and discreteness of the clusters (between sum of squares (SS)) for each inversion.

| Inversion | N outliers | MDS coordinates | Between SS |
|---|---|---|---|
| miinv1.0 | 115 | mds07_neg | 0.954 |
| miinv1.1 | 13 | mds38_pos | 0.956 |
| miinv1.2 | 15 | mds36_pos | 0.868 |
| miinv3.0 | 132 | mds06_pos | 0.985 |
| miinv4.0 | 64 | mds19_pos | 0.939 |
| miinv4.1 | 55 | mds23_neg | 0.962 |
| miinv5.0 | 113 | mds16_neg | 0.955 |
| miinv6.0 | 9 | mds27_neg | 0.959 |
| miinv7.0 | 84 | mds39_pos | 0.883 |
| miinv8.0 | 142 | mds06_neg | 0.962 |
| miinv9.0 | 72 | mds15_pos | 0.978 |
| miinv11.0 | 6 | mds17_neg | 0.958 |
| miinv13.0 | 12 | mds37_pos | 0.847 |
| miinv14.0 | 78 | mds26_pos | 0.888 |
| miinv17.0 | 47 | mds32_neg | 0.879 |
| miinv17.1 | 6 | mds29_neg | 0.933 |
| miinv20.0 | 60 | mds12_pos | 0.852 |

**Table S4.** Trait associations for each inversion using a linear model (*P ≤ 0.05, **P ≤ 0.01, ***P ≤ 0.001). The five mango traits: fruit blush color (BC), fruit firmness (FF), fruit weight (FW), total soluble solids (TSS) and trunk circumference (TC).

| Trait | Inversion | Estimate | Std. Error | t value | Pr(>\|t\|) | Sig. |
|---|---|---|---|---|---|---|
| BC | (Intercept) | 43.56 | 5.19 | 8.38 | 1.21E-14 | |
| BC | miinv1.0 | 4.19 | 1.53 | 2.74 | 6.84E-03 | ** |
| BC | miinv1.1 | 2.12 | 1.35 | 1.56 | 1.19E-01 | |
| BC | miinv1.2 | -1.64 | 1.26 | -1.30 | 1.94E-01 | |
| BC | miinv3.0 | -4.30 | 1.78 | -2.42 | 1.65E-02 | * |
| BC | miinv4.0 | -0.57 | 1.38 | -0.41 | 6.82E-01 | |
| BC | miinv4.1 | -0.85 | 1.63 | -0.52 | 6.02E-01 | |
| BC | miinv5.0 | 0.37 | 1.38 | 0.27 | 7.88E-01 | |
| BC | miinv6.0 | 6.84 | 1.45 | 4.72 | 4.66E-06 | *** |
| BC | miinv7.0 | 1.51 | 1.39 | 1.09 | 2.78E-01 | |
| BC | miinv8.0 | 2.78 | 1.67 | 1.67 | 9.75E-02 | |
| BC | miinv9.0 | 4.76 | 2.01 | 2.37 | 1.89E-02 | * |
| BC | miinv11.0 | -3.60 | 1.83 | -1.97 | 5.08E-02 | |
| BC | miinv13.0 | 0.94 | 1.94 | 0.49 | 6.27E-01 | |
| BC | miinv14.0 | 0.26 | 1.51 | 0.17 | 8.65E-01 | |
| BC | miinv17.0 | 3.40 | 2.28 | 1.49 | 1.38E-01 | |
| BC | miinv17.1 | -2.00 | 1.74 | -1.15 | 2.53E-01 | |
| BC | miinv20.0 | 0.54 | 1.97 | 0.28 | 7.83E-01 | |
| FF | (Intercept) | 1.33 | 0.11 | 11.79 | 3.55E-24 | |
| FF | miinv1.0 | 0.00 | 0.03 | 0.08 | 9.36E-01 | |
| FF | miinv1.1 | -0.03 | 0.03 | -1.09 | 2.77E-01 | |
| FF | miinv1.2 | -0.01 | 0.03 | -0.30 | 7.63E-01 | |
| FF | miinv3.0 | 0.02 | 0.04 | 0.41 | 6.83E-01 | |
| FF | miinv4.0 | -0.03 | 0.03 | -1.08 | 2.84E-01 | |
| FF | miinv4.1 | -0.12 | 0.04 | -3.11 | 2.17E-03 | ** |
| FF | miinv5.0 | -0.04 | 0.03 | -1.41 | 1.61E-01 | |
| FF | miinv6.0 | -0.01 | 0.03 | -0.19 | 8.48E-01 | |
| FF | miinv7.0 | 0.00 | 0.03 | 0.13 | 8.97E-01 | |
| FF | miinv8.0 | -0.06 | 0.04 | -1.61 | 1.09E-01 | |
| FF | miinv9.0 | 0.01 | 0.04 | 0.22 | 8.23E-01 | |
| FF | miinv11.0 | 0.00 | 0.04 | -0.07 | 9.41E-01 | |
| FF | miinv13.0 | -0.06 | 0.04 | -1.43 | 1.54E-01 | |
| FF | miinv14.0 | 0.01 | 0.03 | 0.29 | 7.75E-01 | |
| FF | miinv17.0 | 0.13 | 0.05 | 2.52 | 1.27E-02 | * |
| FF | miinv17.1 | -0.04 | 0.04 | -0.95 | 3.45E-01 | |
| FF | miinv20.0 | 0.11 | 0.04 | 2.51 | 1.31E-02 | * |
| FW | (Intercept) | 19.61 | 1.58 | 12.39 | 6.73E-26 | |
| FW | miinv1.0 | -0.79 | 0.47 | -1.67 | 9.60E-02 | |
| FW | miinv1.1 | 0.27 | 0.41 | 0.67 | 5.04E-01 | |
| FW | miinv1.2 | 0.21 | 0.39 | 0.54 | 5.91E-01 | |
| FW | miinv3.0 | 0.09 | 0.55 | 0.16 | 8.69E-01 | |
| FW | miinv4.0 | -0.22 | 0.43 | -0.50 | 6.18E-01 | |
| FW | miinv4.1 | 0.03 | 0.55 | 0.06 | 9.53E-01 | |
| FW | miinv5.0 | 0.28 | 0.42 | 0.66 | 5.13E-01 | |
| FW | miinv6.0 | -2.16 | 0.45 | -4.79 | 3.49E-06 | *** |

| Trait | Inversion | Estimate | Std. Error | t value | Pr(>\|t\|) | Sig. |
|---|---|---|---|---|---|---|
| FW | miinv7.0 | 1.18 | 0.44 | 2.70 | 7.51E-03 | ** |
| FW | miinv8.0 | 0.93 | 0.52 | 1.80 | 7.33E-02 | |
| FW | miinv9.0 | -1.07 | 0.61 | -1.77 | 7.92E-02 | |
| FW | miinv11.0 | 0.33 | 0.59 | 0.56 | 5.79E-01 | |
| FW | miinv13.0 | -0.64 | 0.58 | -1.09 | 2.76E-01 | |
| FW | miinv14.0 | -0.21 | 0.46 | -0.46 | 6.43E-01 | |
| FW | miinv17.0 | -0.59 | 0.70 | -0.84 | 3.99E-01 | |
| FW | miinv17.1 | 0.19 | 0.55 | 0.35 | 7.27E-01 | |
| FW | miinv20.0 | -2.35 | 0.61 | -3.87 | 1.53E-04 | *** |
| TSS | (Intercept) | 1.20 | 0.04 | 33.61 | 1.69E-79 | |
| TSS | miinv1.0 | -0.01 | 0.01 | -1.00 | 3.18E-01 | |
| TSS | miinv1.1 | -0.02 | 0.01 | -2.08 | 3.89E-02 | * |
| TSS | miinv1.2 | -0.01 | 0.01 | -0.98 | 3.28E-01 | |
| TSS | miinv3.0 | -0.02 | 0.01 | -1.59 | 1.13E-01 | |
| TSS | miinv4.0 | 0.02 | 0.01 | 1.80 | 7.31E-02 | |
| TSS | miinv4.1 | -0.03 | 0.01 | -2.74 | 6.85E-03 | ** |
| TSS | miinv5.0 | 0.01 | 0.01 | 1.29 | 1.97E-01 | |
| TSS | miinv6.0 | 0.00 | 0.01 | -0.39 | 6.96E-01 | |
| TSS | miinv7.0 | -0.01 | 0.01 | -1.47 | 1.43E-01 | |
| TSS | miinv8.0 | 0.01 | 0.01 | 0.91 | 3.64E-01 | |
| TSS | miinv9.0 | 0.02 | 0.01 | 1.55 | 1.22E-01 | |
| TSS | miinv11.0 | -0.01 | 0.01 | -0.89 | 3.74E-01 | |
| TSS | miinv13.0 | 0.00 | 0.01 | -0.12 | 9.05E-01 | |
| TSS | miinv14.0 | 0.00 | 0.01 | 0.21 | 8.37E-01 | |
| TSS | miinv17.0 | 0.02 | 0.02 | 1.31 | 1.92E-01 | |
| TSS | miinv17.1 | 0.00 | 0.01 | -0.36 | 7.21E-01 | |
| TSS | miinv20.0 | 0.00 | 0.01 | -0.35 | 7.30E-01 | |
| TC | (Intercept) | 55.24 | 4.75 | 11.62 | 5.18E-23 | |
| TC | miinv1.0 | -1.15 | 1.34 | -0.86 | 3.92E-01 | |
| TC | miinv1.1 | 1.24 | 1.18 | 1.05 | 2.95E-01 | |
| TC | miinv1.2 | -0.39 | 1.08 | -0.36 | 7.18E-01 | |
| TC | miinv3.0 | -0.14 | 1.68 | -0.09 | 9.32E-01 | |
| TC | miinv4.0 | -1.64 | 1.21 | -1.35 | 1.78E-01 | |
| TC | miinv4.1 | 0.19 | 1.61 | 0.12 | 9.07E-01 | |
| TC | miinv5.0 | -0.13 | 1.21 | -0.11 | 9.12E-01 | |
| TC | miinv6.0 | 0.85 | 1.28 | 0.67 | 5.05E-01 | |
| TC | miinv7.0 | 0.56 | 1.25 | 0.45 | 6.54E-01 | |
| TC | miinv8.0 | 1.40 | 1.51 | 0.92 | 3.56E-01 | |
| TC | miinv9.0 | 1.73 | 1.77 | 0.97 | 3.31E-01 | |
| TC | miinv11.0 | -1.18 | 1.62 | -0.73 | 4.66E-01 | |
| TC | miinv13.0 | -1.91 | 1.68 | -1.14 | 2.56E-01 | |
| TC | miinv14.0 | 0.18 | 1.33 | 0.13 | 8.94E-01 | |
| TC | miinv17.0 | -9.72 | 2.02 | -4.82 | 3.34E-06 | *** |
| TC | miinv17.1 | 2.76 | 1.72 | 1.60 | 1.11E-01 | |
| TC | miinv20.0 | 1.18 | 1.82 | 0.65 | 5.19E-01 | |

**Table S5.** Trait associations for each inversion using the iterative random forest importance scores. The five mango traits: fruit blush color (BC), fruit firmness (FF), fruit weight (FW), total soluble solids (TSS) and trunk circumference (TC). The combined variance explained for all inversions is shown per trait.

| Inv | BC | FF | FW | TC | TSS |
|---|---|---|---|---|---|
| miinv6.0 | 6941 | 0.52 | 346 | 463 | 0.15 |
| miinv1.1 | 3677 | 0.74 | 261 | 1352 | 0.10 |
| miinv7.0 | 486 | 0.50 | 244 | 231 | 0.10 |
| miinv13.0 | 277 | 0.53 | 240 | 497 | 0.02 |
| miinv1.0 | 966 | 0.52 | 223 | 315 | 0.07 |
| miinv8.0 | 1258 | 0.46 | 197 | 628 | 0.02 |
| miinv4.0 | 2310 | 0.67 | 154 | 1345 | 0.11 |
| miinv4.1 | 2850 | 0.78 | 130 | 318 | 0.15 |
| miinv17.0 | 1105 | 0.50 | 105 | 3586 | 0.02 |
| miinv1.2 | 694 | 0.55 | 104 | 638 | 0.06 |
| miinv20.0 | 268 | 0.47 | 100 | 147 | 0.02 |
| miinv17.1 | 1347 | 0.65 | 98 | 588 | 0.04 |
| miinv5.0 | 1807 | 0.77 | 95 | 1753 | 0.12 |
| miinv9.0 | 4043 | 0.58 | 87 | 71 | 0.06 |
| miinv11.0 | 879 | 0.38 | 78 | 1516 | 0.01 |
| miinv14.0 | 2397 | 0.58 | 59 | 440 | 0.01 |
| miinv3.0 | 2748 | 0.51 | 23 | 473 | 0.16 |
| **var. explained** | 0.33 | 0.02 | 0.12 | 0.16 | 0.21 |

**Table S6.** The number and percentage of mutations for each substitution type and SIFT prediction category. The three substitution types include nonsynonymous (protein altering), synonymous (nonprotein altering), and start-lost (nonsense mutations). The SIFT prediction categories include deleterious (SIFT < 0.05; mutation in conserved region) and tolerated (SIFT > 0.05; mutation in non-conserved region).

| **Substitution SIFT prediction** | **N mutations** | **% of Total** |
|---|---|---|
| Nonsynonymous deleterious | 36,606 | 18.6 |
| Nonsynonymous tolerated | 79,002 | 40.2 |
| Synonymous deleterious | 830 | 0.4 |
| Synonymous tolerated | 80,162 | 40.8 |
| Start-lost deleterious | 67 | <0.01 |
| Start-lost tolerated | 51 | <0.01 |
| Total | 196,718 | |

**Table S7.** The 11 QTL regions and their mean deleterious score. These inversions have a deleterious score lower (-) or higher (+) than the genome average (quantiles 2.5% = 0.517 and 97.5% = 0.513 from 1,000 bootstraps). Deleterious scores are represented as + >0.487, ++ >0.524, +++ >0.560, - <0.483, - - <0.443 and, - - - <0.402. No deleterious score for miTCQTL12 as it is in a missing/non-coding region.

| QTL | N GWAS hits | QTL size (kb) | Start (Mb) | End (Mb) | Deleterious score | Deleterious score range | Genes within QTL |
|---|---|---|---|---|---|---|---|
| miBCQTL15 | 44 | 136 | 10.65 | 10.78 | 0.320 | - - - | 4 |
| miFFQTL5 | 7 | 89 | 11.96 | 12.05 | 0.447 | - | 13 |
| miFFQTL9 | 7 | 217 | 8.89 | 9.10 | 0.505 | + | 15 |
| miTCQTL1 | 9 | 75 | 22.65 | 22.72 | 0.369 | - - - | 8 |
| miTCQTL2 | 7 | 3032 | 20.69 | 23.72 | 0.494 | + | 446 |
| miTCQTL6 | 7 | 34 | 18.45 | 18.49 | 0.574 | +++ | 5 |
| miTCQTL14 | 7 | 51 | 3.91 | 3.96 | 0.393 | - - | 11 |
| miTSSQTL5 | 67 | 3185 | 6.44 | 9.62 | 0.482 | - | 171 |
| miFWQTL11 | 10 | 18 | 5.72 | 5.74 | 0.411 | - - | 3 |
| miFWQTL18 | 6 | 515 | 10.74 | 11.25 | 0.473 | - | 59 |
| miTCQTL12 | 4 | 16 | 4.08 | 4.09 | Noncoding region and missing region | | 1 |
